# Tiled base editing screens uncover rare second-site mutations that rescue a misfolded disease protein

**DOI:** 10.64898/2026.10.30.763201

**Authors:** Shivali Rawat, Parijat Sarkar, Eric Giavedoni, Ganesh V. Pusapati, Murugesh Padmanarayana, Mallory Shin, Elizabeth Phillippi, Taylar Simmons, Ruth Azaria, Diwakar Shukla, Mark L. Schultz, Rajat Rohatgi

**Author notes:** These authors contributed equally (Shivali Rawat, Parijat Sarkar).

## Abstract

Therapeutic base editing can directly revert ∼30% of disease-causing missense mutations to the wild-type sequence. One strategy to broaden its reach is to install second-site corrector mutations that improve protein stability or function across multiple pathogenic alleles. Here, we use tiled adenine and cytosine base editing screens to identify such corrector mutations in NPC1, a lysosomal cholesterol transporter whose misfolding causes Niemann-Pick type C disease. Tiling mutagenesis of the endogenous *NPC1* locus in haploid human cells carrying the prevalent *NPC1^I1061T^* disease allele, combined with phenotypic reporters for cholesterol transport and protein abundance, uncovered rare intragenic suppressor mutations distributed across multiple domains. Unexpectedly, we identified a promiscuous second-site variant that can correct many different pathogenic *NPC1* mutations. These correctors improve protein stability, promote ER exit and lysosomal trafficking, and restore cholesterol clearance in patient-derived fibroblasts and induced neurons. Our results establish *in situ* base editing screens for the discovery of second-site corrector mutations that are actionable for therapeutic base editing when the original mutation cannot be corrected.

## Introduction

Some diseases are caused by missense mutations that disrupt folding of the encoded protein but often do not abolish its intrinsic biochemical activity. A prominent example is the ΔF508 mutation in the cystic fibrosis transmembrane conductance regulator (CFTR), which causes the majority of cystic fibrosis (CF) cases. A critical early insight came from experiments showing that reduced temperature (*1*) or second-site suppressor mutations (*2*) could partially rescue CFTR-ΔF508 function by allowing some of the mutant protein to escape endoplasmic reticulum (ER) quality control and reach the plasma membrane. This insight spurred development of pharmacological chaperones, called “correctors”, that have transformed the treatment of CF(*3*, *4*).

The clinical success of CFTR folding correctors suggests that a similar strategy could be applied to the treatment of other diseases caused by mutations in ER-transiting membrane proteins. Niemann-Pick type C disease is a devastating pediatric neurodegenerative disorder caused by mutations in the genes encoding Niemann-Pick C1 (NPC1) and C2 (NPC2), two proteins that cooperate to export cholesterol from the lysosome (**Fig.1a**). NPC2, a soluble luminal protein, binds unesterified cholesterol released from low-density lipoprotein (LDL) derived cholesterol esters and transfers it to NPC1, which facilitates cholesterol transport across the lysosomal membrane to other cellular destinations including the ER and plasma membrane. Loss of NPC1 or NPC2 function causes accumulation of cholesterol and sphingolipids in lysosomes, leading to lysosomal dysfunction and progressive neurodegeneration.

**Figure 1:**
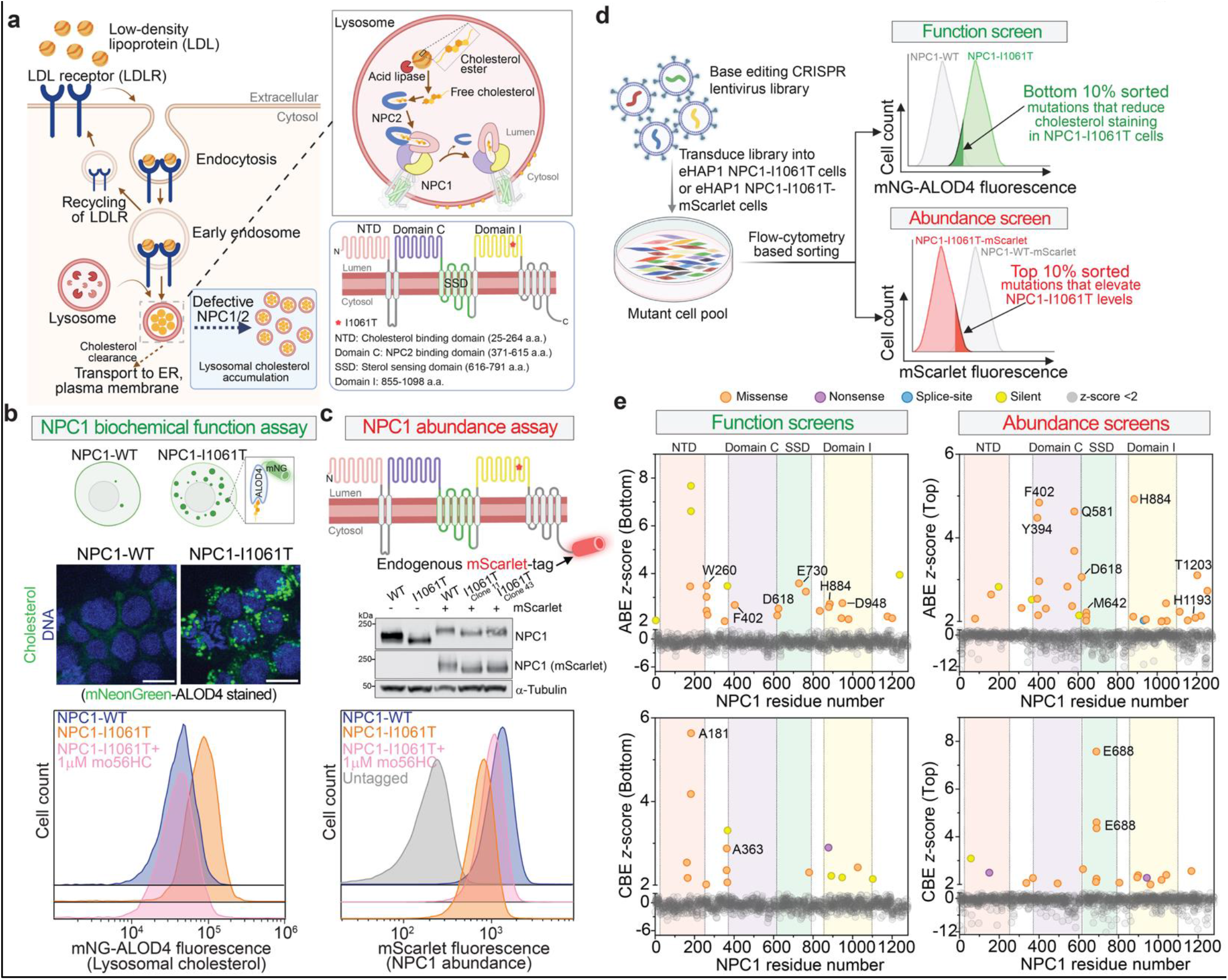
Tiled base editing screens for mutations that rescue NPC1-I1061T function and abundance. **a**, Low-density lipoprotein (LDL) is endocytosed and trafficked to lysosomes, where free cholesterol is liberated by acid lipase and exported by NPC2 and NPC1. The prevalent misfolding mutation I1061T (red star) lies in domain I of NPC1. Domain boundaries and functions are indicated. **b,** Lysosomal cholesterol measured by mNeonGreen-ALOD4 (mNG-ALOD4) staining (top). Confocal images (middle) and flow cytometry histograms (bottom) of mNG-ALOD4–stained NPC1-WT and NPC1-I1061T eHAP1 cells (Scale bar: 10 µm). **c,** Endogenous C-terminal mScarlet tagging of NPC1 (top). Immunoblot (middle) and flow cytometry histograms (bottom) show abundance of NPC1-WT and NPC1-I1061T. NPC1-I1061T cells show elevated mNG-ALOD4 signal (**b**) and reduced mScarlet (**c**) versus WT; both are partially restored by the pharmacological chaperone mo56HC (1 µM, 48 hr). **d,** Pooled base editing screen strategy. Lentiviral libraries co-encoding ABE or CBE with respective sgRNAs tiling all *NPC1* exons, were transduced into NPC1-I1061T cells (function screen) or NPC1-I1061T-mScarlet cells (abundance screen). The bottom 10% by mNG-ALOD4 (function rescue) or top 10% by mScarlet (abundance rescue) were isolated by FACS. **e,** Plots showing sgRNA z-scores for ABE (top panel) and CBE (bottom panel) function (left) and abundance (right) screens versus NPC1 residue number, computed against a non-targeting control distribution (n=2). For sgRNAs predicted to generate multiple mutations, the first residue in the editing window is plotted. Dots colored by predicted mutation type (orange, missense; purple, nonsense; blue, splice-site; yellow, silent); z-score <2 in gray. Domain shading as in **a**. See also **figs. S1,2 and Supplementary Tables 1 and 2**.

Missense mutations in *NPC1* cause >95% of Niemann-Pick type C disease cases (hereafter called Niemann-Pick type C1 disease). Many disrupt NPC1 folding, resulting in ER retention and degradation by ER-associated degradation (ERAD) and ER-phagy pathways (*5–7*). For the prevalent *NPC1* I1061T mutation, the protein retains intrinsic biochemical activity, but too little reaches the lysosome. As with CFTR-ΔF508, increasing lysosomal abundance of NPC1-I1061T through overexpression, reduced temperature, or pharmacological chaperones like mo56-hydroxycholesterol (mo56HC) partially rescues cholesterol transport (*5*, *8–11*). However, clinically viable corrector drugs against folding and trafficking defective mutants of NPC1 have yet to be developed.

The identification of second-site suppressor mutations can provide uniquely valuable information about the misfolding and mis-trafficking mechanisms, conformational states, and druggability of mutant, disease-causing proteins. Suppressor mutations identify residues and structural regions of the protein that can be targeted by small molecules or genome editing approaches to restore protein function. However, isolation of such (expectedly rare) mutations remains challenging in mammalian cells. Deep mutational scanning (DMS) is the most comprehensive way to identify such mutants but cannot be easily applied to large proteins like NPC1 and requires overexpression from a non-native genomic locus (*12*). Base editing has emerged as an alternative scanning mutagenesis method that allows the introduction of transition mutations across endogenous genomic loci using Cas9-derived editors and single guide RNAs (sgRNAs) (*13–18*). While base editing samples only a fraction of the mutational landscape, it can be easily implemented using standard pooled CRISPR screening methods coupled with phenotypic selection and thus enables the assessment of mutations in endogenous proteins in living cells. Importantly, every corrector mutation discovered in a base editing screen can, in principle, be therapeutically installed by the same editor.

Motivated by these advantages, we asked if base editing screens could be used to identify intragenic suppressor mutations in *NPC1^I1061T^* in human cells, analogous to how screens in yeast initially identified corrector mutations in CFTR-ΔF508 (*2*). Despite sampling only a fraction of the NPC1 sequence landscape, we identified corrector mutations at distant sites that improve NPC1-I1061T protein stability, promote ER exit and lysosome trafficking, and restore cholesterol transport. Interestingly, we also identified mutations that selectively impact protein stability or trafficking, without improving cholesterol transport activity. Thus, base editing screens can be used to identify mutations that stabilize different states in the lifecycle of a complex multi-domain membrane protein, providing valuable tools for understanding protein biogenesis, function, and structure. Our work provides a general framework for using base editing screens to discover therapeutically installable corrector mutations for protein misfolding diseases.

## Results

### Screens to identify mutations that improve protein function and abundance

Human haploid cells (eHAP1) provide an ideal model for base editing screens since they only possess one allele of each gene, allowing the phenotypic evaluation of recessive mutations (*18–21*). eHAP1 cells have been used to model the cellular defects in Niemann-Pick type C1 disease, and base editing in eHAP1 cells allows the functional evaluation of *NPC1* disease variants (*22*). We used genome editing to install the I1061T mutation at the single *NPC1* locus in multiple independent clonal eHAP1 cell lines and confirmed biochemical and molecular Niemann-Pick type C1 disease phenotypes using a panel of assays (**Fig.1b, fig. S1a-d**). NPC1-I1061T cells demonstrated massive cholesterol accumulation in lysosomes after an LDL cholesterol pulse (**Fig.1b**). A cholesterol binding domain from a bacterial toxin fused to a fluorescent protein (mNeonGreen-tagged ALOD4 or mNG-ALOD4, **fig. S1e**) allowed us to both visualize (by fluorescence microscopy) and measure (by flow cytometry) lysosomal cholesterol in individual cells (*23*) (**Fig.1b**). To measure NPC1-I1061T protein abundance in individual eHAP1 cells by flow cytometry, we introduced an in-frame mScarlet tag at the C-terminus of the single *NPC1* allele (**Fig.1c, fig. S1f-h**). NPC1-I1061T abundance was reduced compared to wild-type (WT) NPC1 and it migrated faster in SDS-PAGE, consistent with ER trapping (*5*, *24*) (**Fig.1c**). In a control experiment, the pharmacological chaperone mo56HC increased NPC1-I1061T protein abundance and concomitantly reduced lysosomal cholesterol (*8*) (**Fig.1b,c**). In summary, our assays and engineered eHAP1 cell lines enabled us to measure lysosomal cholesterol, a measure of NPC1 biochemical activity, and NPC1 protein abundance in single cells, thus enabling the isolation of rare cells with desired activity or abundance phenotypes from a pooled mutant library by fluorescence-activated cell sorting (FACS).

To generate pooled mutant libraries in NPC1-I1061T and NPC1-I1061T-mScarlet eHAP1 cells, we used promiscuous PAM-flexible (NG-Cas9n) adenine and cytosine base editors (NG-ABE8e and NG-BE3.9) to introduce transition mutations across all *NPC1* exons (*13*, *14*, *25*, *26*). Our ABE and CBE libraries are predicted to generate missense mutations at 80-90% of amino acid residues encoded by *NPC1* (**Supplementary Tables 1 and 2**). Both libraries also generated silent and splice-site mutations, since the design strategy allowed sgRNAs to extend ∼20 nucleotides into introns at each intron-exon junction (**fig. S1i**). However, nonsense mutations can only be generated by the CBE editor. The sgRNA library covers each exon in a tiled fashion, such that many codons are targeted by multiple sgRNAs (**fig. S1j**).

Separate screens for second-site mutations that improve NPC1-I1061T function and abundance were conducted in NPC1-I1061T and NPC1-I1061T-mScarlet eHAP1 cells, respectively (**Fig.1d**). To isolate mutant cells with improved lysosomal cholesterol transport, we used FACS to sort cells in the bottom 10% of the mNG-ALOD4 fluorescence intensity distribution. Conversely, mutant cells with improved protein abundance were isolated based on mScarlet fluorescence values that were in the top 10% of the overall distribution of the mutant population. Illumina sequencing was used to measure sgRNA frequencies in these phenotypically sorted populations and matched parent unsorted populations (**fig. S1k**). We used two methods to analyze the data from these screens. First, enrichment for each sgRNA was calculated as the log_2_ fold change (LFC) between each sorted population and its matched unsorted population. The distribution of LFCs from 278 non-targeting control (NTC) guides was subsequently used to standardize these values and derive z-scores (**Supplementary Tables 1 and 2, fig. S2a**). Second, screens were independently analyzed using MAGeCK, a widely-used, open-source analysis program (*27*, *28*) (**Supplementary Tables 1 and 2**).

A z-score threshold of ≥2 was used as a cutoff for filtering candidate sgRNAs for subsequent analyses. Manhattan-type z-score plots (**Fig.1e**), lollipop plots (**fig. S2b**), and z-score distributions (**fig. S2c**) all highlighted the rarity of corrector mutations. Cross-correlation analysis of z-scores for each sgRNA between the screens pointed to sgRNAs that increased both abundance and function of NPC1-I1061T, a phenotype expected of a mutation that improves folding, but also to sgRNAs that increased abundance without improving function (**fig. S2d, Fig.2a**). Hence, paired abundance and function screens allowed the identification of second-site mutations that impact different cell biological behaviors of NPC1-I1061T, likely reflecting the ability of these mutations to stabilize different conformations of this multi-domain transporter. Notably, corrector mutations were not confined to the luminal domain that contains I1061T (domain I), but were distributed across luminal, transmembrane and even cytoplasmic-facing regions of NPC1-I1061T (**Fig.2a**).

### Identification of specific mutations that improve NPC1-I1061T function and abundance

To select mutations for mechanistic analysis, we followed a systematic filtering and validation strategy (diagrammed in **fig. S3a**). sgRNAs with z-scores ≥2 (predicted mutations mapped onto the NPC1 structure in **Fig.2a**) from the function and abundance screens were individually introduced into eHAP1 NPC1-I1061T and NPC1-I1061T-mScarlet cells, respectively, followed by measurement of lysosomal cholesterol and NPC1 protein abundance (**Fig.2b**). We chose to further characterize sgRNAs with the largest effects in our validation studies: four that improved both cholesterol transport and protein abundance and two that only improved abundance but not function (**Fig.2b-d**). Interestingly, these sgRNAs are predicted to generate mutations (**Fig.2c**) proximal to a central tunnel in NPC1 that may serve as a conduit for cholesterol transport (**Fig.2e**): the Cα atoms of 6 out of 7 mutations made by these sgRNAs are within 10 Å of the tunnel surface. Cholesterol is thought to enter the tunnel at its luminal end (from the NTD), traverse through the protein, and ultimately move across a gate (**Fig.2e**, inset) between the membrane-proximal end of the tunnel and the SSD to insert into the lysosomal membrane (*29–32*). Base editors can make mutations at multiple codons within a window spanning N3-N10 of the protospacer. Consequently, introduction of an sgRNA and base editor produces an oligoclonal population of cells carrying various combinations of mutations, only some of which cause the phenotype of interest (**fig. S3b**). To parse which amino acid changes led to improvements in function and abundance, we prospectively isolated clonal cell lines after transfecting each of the sgRNAs chosen for follow-up into eHAP1 NPC1-I1061T-mScarlet cells (**Fig.2c**). The specific second-site mutation present in each clonal cell line was determined by amplicon sequencing (**fig. S3c**). As a control, we used an sgRNA targeting the parent I1061T mutation. Clones in which the sequence was changed back to WT (T1061I) showed both improved protein abundance and reduced lysosomal cholesterol accumulation (**Fig.2f,g**). However, a clone carrying a different mutation (T1061S) at the same residue failed to show correction of either phenotype. For each of the other sgRNAs, correction of the abundance and cholesterol transport phenotypes also required specific amino acid changes. For example, the double mutation F402S-F403P improved the cholesterol transport activity and abundance of NPC1-I1061T, but clones with related mutations (F402S-F403S, F403P) did not show rescue (**Fig.2f,g**). Based on this clonal analysis, we identified F402S-F403P (A5), D618G-S619G (A6), H884R (A12) and H1193R (A20) as second-site correctors that improve both protein abundance and activity of NPC1-I1061T (**Fig.2f,g**). E688K (C29) and Y394C (A35) mutations increased NPC1-I1061T abundance, but did not improve cholesterol transport activity (**Fig.2f,g**).

We also introduced sgRNAs encoding these mutations into eHAP1 cells with WT NPC1-mScarlet (**fig. S4a**). All four second-site correctors had no effect on the activity or abundance of WT NPC1 (**fig. S4b-d)**; however, E688K and Y394C abolished cholesterol transport activity (**fig. S4c,d)**. Taken together, these data suggest that second-site corrector mutations require very specific changes in the amino acid side chains. Additionally, such mutations can separate protein abundance from cholesterol transport activity, presumably by stabilizing an inactive conformation of either WT NPC1 or NPC1-I1061T.

### Corrector mutations rescue NPC1 abundance, trafficking, localization, and function

To evaluate the effect of second-site corrector mutations on various properties of NPC1-I1061T-mScarlet, we compared them to the T1061I reversion installed using the same base editing and clonal isolation pipeline (**Fig.2f,g**). Each of the second-site mutations increased steady-state protein abundance of NPC1-I1061T, with H1193R (on the cytoplasmic surface) having the smallest effect (**Fig.3a**). Both mutations that failed to rescue function (E688K and Y394C) elevated NPC1-I1061T protein abundance to levels higher than WT NPC1. Cycloheximide chase experiments demonstrated that all second-site correctors lengthened the half-life of NPC1-I1061T (**Fig.3b, fig. S5a**).

To measure NPC1-I1061T trafficking from the ER to the Golgi, we tested the sensitivity to the glycosidases Endo H and PNGase F (*6*) (**Fig.3c**). Endo H removes N-glycans added to NPC1 in the ER, causing an increase in its mobility by SDS-PAGE. Properly folded NPC1 molecules that pass the ER Quality Control (ERQC) filter move to the medial Golgi, where the attached N-glycans are further modified such that they become resistant to Endo H (but remain sensitive to PNGase F). Hence, the fraction of NPC1 resistant to Endo H is a measure of the fraction of properly folded protein that has passed the ERQC system and trafficked to the Golgi. The four mutations that rescued NPC1-I1061T function increased the fraction of NPC1-I1061T resistant to Endo H (**Fig.3c,d**). Of the two second-site mutations that only improved abundance, E688K increased the Endo H resistant fraction (consistent with an improvement in folding). Unexpectedly, NPC1-I1061T-Y394C remained fully Endo H sensitive (**Fig.3c,d**), suggesting it was trapped in the ER despite being more stable than NPC1-I1061T (**Fig.3b**). Colocalization studies using immunofluorescence (IF) demonstrated that all second-site corrector mutations except for Y394C increased the abundance of NPC1-I1061T in late endosomes and lysosomes, compartments where NPC1 must be localized to transport cholesterol (**Fig.3e, fig. S5b**).

Four corrector mutations that improved trafficking to the lysosome also increased cholesterol clearance from lysosomes after an LDL pulse, measured by staining either with mNG-ALOD4 (**Fig.3f,g**) or filipin, a polyene macrolide antibiotic that binds cholesterol (*33*) (**fig. S5c,d**). Membrane dyshomeostasis caused by loss of NPC1 activity leads to increased lysosome fragility, which can be assessed by sensitivity to the membrane destabilizing amyloidogenic agent L-leucyl-L-leucine methyl ester (LLOMe) (*34*, *35*). The four second-site corrector mutations that increased NPC1-I1061T trafficking to the lysosome and reduced cholesterol accumulation also increased resilience to LLOMe, consistent with an improvement in lysosomal membrane integrity (**Fig.3h, fig. S5e**). NPC1-I1061T-Y394C, which remained trapped in the ER, failed to rescue cholesterol transport or reduce sensitivity to LLOMe. NPC1-I1061T-E688K also did not improve either metric of NPC1 function despite being localized correctly in the lysosome (**Fig.3e**), likely because it cannot transport cholesterol. The impact of the E688K mutation in NPC1 resembles that of itraconazole and posaconazole, antifungals that inhibit NPC1 function but improve its folding and trafficking to the lysosome (*36*). Washout of these inhibitors after a treatment pulse improves lysosomal cholesterol clearance in NPC1-I1061T cells (*11*).

### Molecular dynamics simulations highlight a role for the sterol-sensing domain in NPC1 stability and activity

To understand how second-site correctors improve folding of NPC1-I1061T, we performed all-atom molecular dynamics (MD) simulations of full-length NPC1. Though NPC1 is a lysosomal protein, we chose to conduct these simulations under conditions that mimic the ER, the environment in which NPC1 folds. We started with an NPC1 structure solved at neutral pH (PDB 6UOX) with the best wwPDB scores, maintained neutral pH during the simulations, and embedded NPC1 variants in a symmetric lipid bilayer mimicking the lipid composition of the ER membrane (*32*, *37*) (**Fig.4a,b, Supplementary Table 4**). Cholesterol or other ligands identified in various NPC1 structures were excluded, since the cholesterol content of the ER is low and it is not clear that NPC1 binds cholesterol during folding.

We performed 30 µs simulations of four systems: WT NPC1, NPC1-I1061T, NPC1-I1061T-D618G-S619G, and NPC1-I1061T-E688K. We selected these variants because D618, S619, and E688 cluster at the luminal-facing border of the sterol-sensing domain (SSD) (**Fig. 2e**, inset), yet produce distinct rescue phenotypes: D618G-S619G rescues both NPC1-I1061T trafficking and cholesterol-transport function, whereas E688K rescues trafficking but not function. This region has also been proposed to form a hydrophilic gate that regulates cholesterol transfer from the central tunnel into the SSD at the luminal leaflet of the membrane (*29*– *32, 38*). Indeed, D618 (located at the luminal border of TM3) and E688 (located at the luminal border of TM5) are conserved in the yeast NPC1 homolog NCR1 and have been implicated in regulating sterol movement through this gate between the central tunnel and SSD (*30*).

**Figure 2:**
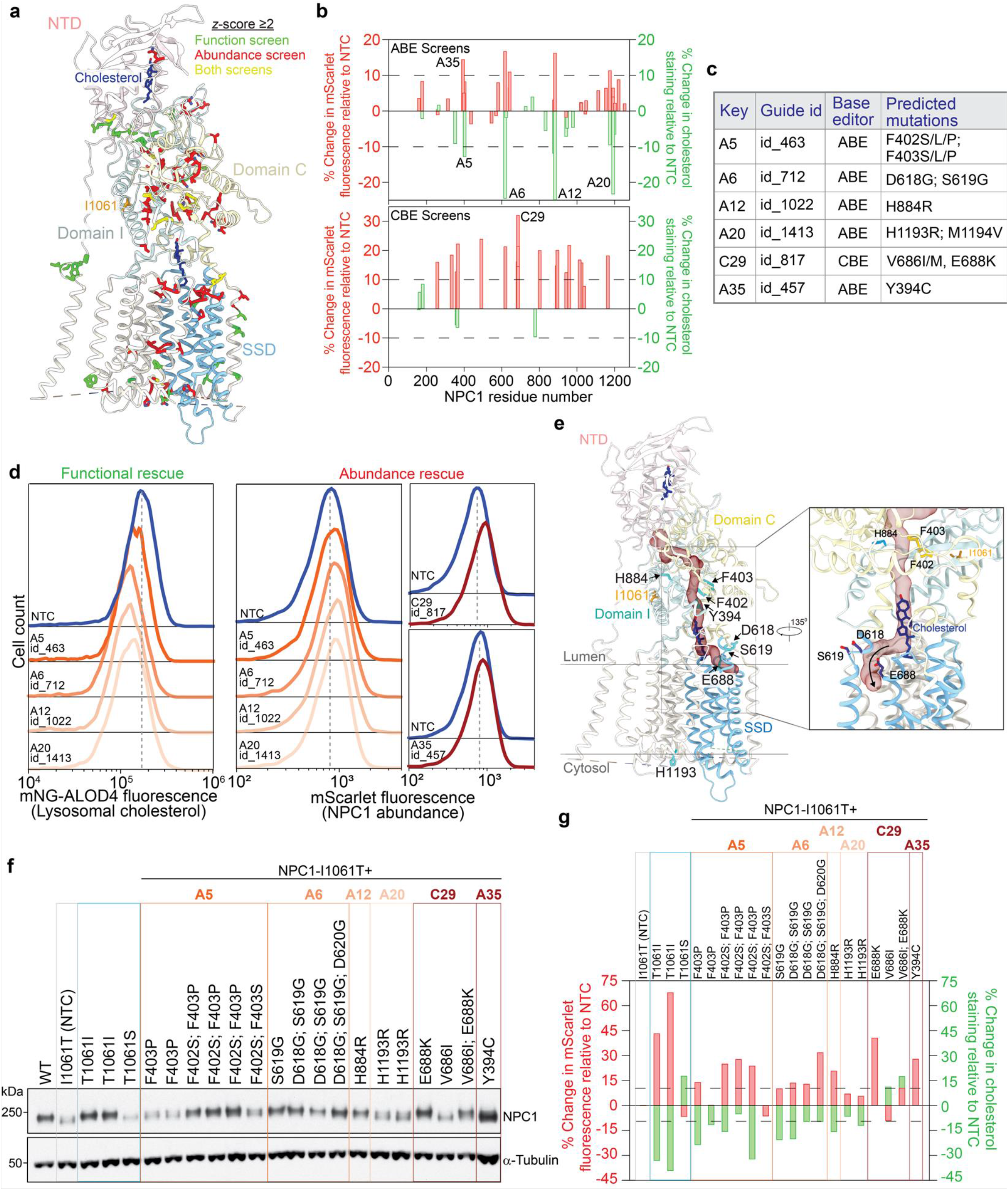
Individual validation and clonal isolation of second-site corrector mutations of NPC1-I1061T. **a**, Residues targeted by sgRNAs with z-score ≥2 (**Fig.1e**) mapped onto the NPC1 structure (PDB: 6W5U(*29*)), colored by enrichment in the function screen (green), abundance screen (red), or both (yellow). **b,** Individual validation of screen hits filtered as shown in **fig. S3a**. Each sgRNA was introduced into eHAP1 NPC1-I1061T (function) or eHAP1 NPC1-I1061T-mScarlet (abundance) cells. Percent change in lysosomal cholesterol (mNG-ALOD4) and NPC1 abundance (mScarlet) relative to I1061T carrying a non-targeting control (NTC) sgRNA was measured by flow cytometry. ABE (top panel) and CBE (bottom panel) hits are plotted separately; labeled bars indicate the six sgRNAs selected for follow-up (n=1). **c,** Summary of the six selected sgRNAs, their base editor, and predicted mutations. **d,** Flow cytometry histograms for the six sgRNAs versus NTC, measuring lysosomal cholesterol (mNG-ALOD4; functional rescue) or NPC1 abundance (mScarlet; abundance rescue). **e,** NPC1 structure with the cholesterol tunnel (MOLE 2.5 (*50*)) shown in transparent surface representation. The six sgRNAs targeted residues are highlighted. Inset: several corrector mutations cluster around a gate between the central tunnel and the SSD. **f,** Immunoblot measuring NPC1 abundance (α-tubulin, loading control) from clonal cell lines isolated from each base-edited population, carrying the indicated single or compound mutations on the NPC1-I1061T background. **g,** Flow cytometry quantification of percent change in cholesterol clearance and NPC1 abundance versus NTC in clonal lines from **f** (n=1). See also **fig. S3 and Supplementary Table 3**.

To identify the dominant conformational states sampled by each variant, we applied time-lagged independent component analysis (tICA), a dimensionality reduction method that extracts the slowest collective motions from simulation trajectories (*39, 40*). The SSD of WT NPC1 predominantly occupied a single conformational state (state α; **Fig.4a**), but the NPC1-I1061T SSD populated a second distinct state (state β) in the conformational landscape that was rarely visited by the wild-type protein. This aberrant state was characterized by a displacement of SSD helices TM3, TM4, and TM5, with TM3 showing the largest inward displacement (RMSD = 5.2 Å) relative to wild-type (**Fig.4b**). Addition of the D618G-S619G corrector mutations to NPC1-I1061T restored a WT-like conformational distribution, largely eliminating the β state population (**Fig.4a**). The E688K mutation also reduced β state occupancy, but failed to completely restore a WT conformational distribution.

**Figure 3:**
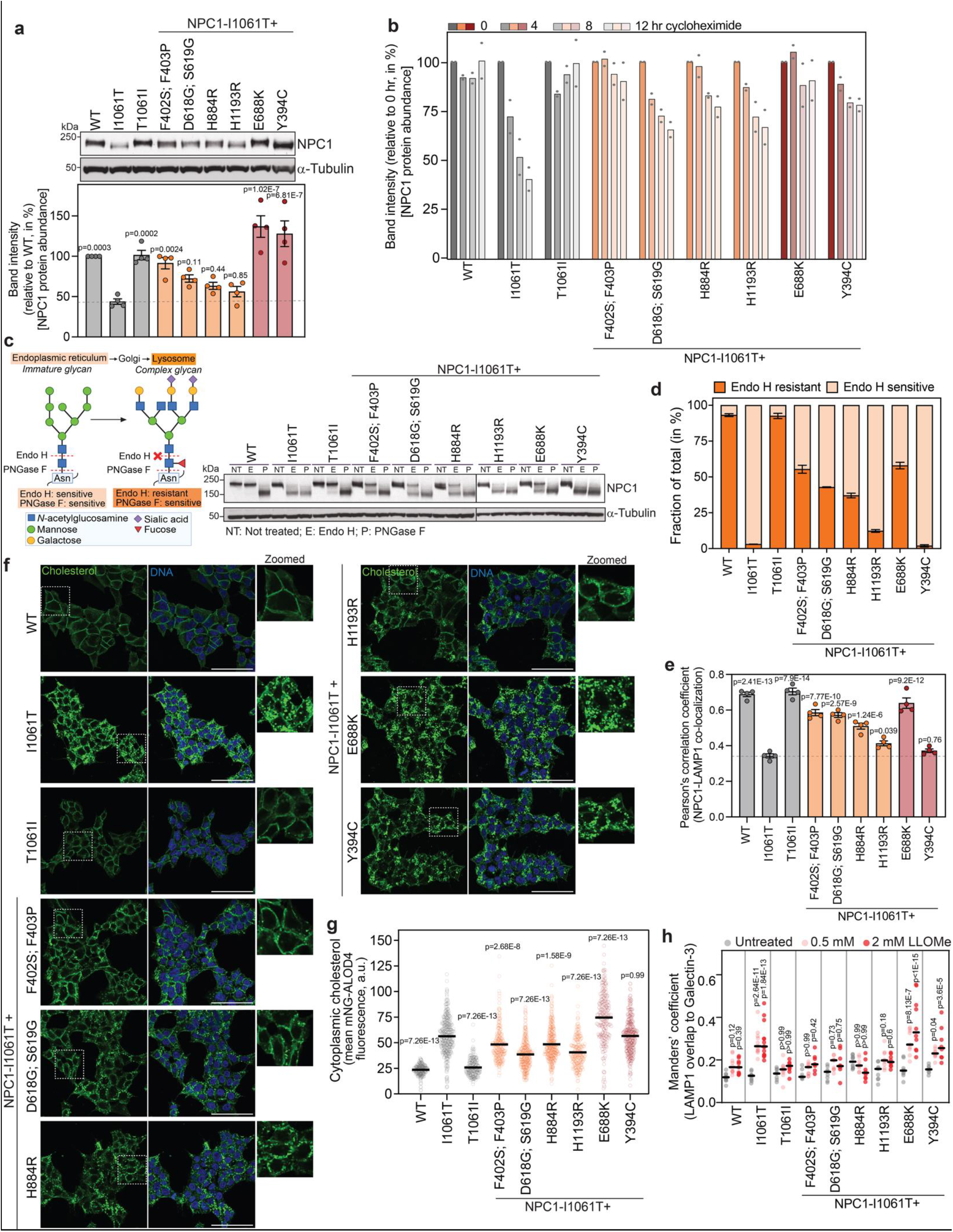
Second-site mutations differentially restore stability, ER exit, lysosomal localization, and cholesterol transport of NPC1-I1061T. **a**, Steady-state NPC1 abundance by immunoblot (top) and quantification (bottom) in clonal eHAP1 lines carrying the indicated mutations on the NPC1-I1061T-mScarlet background; α-tubulin as loading control (n=4, mean ± SEM). **b,** NPC1 half-life by cycloheximide chase, normalized to 0 hr (n=2). **c,d,** ER exit assayed by Endo H and PNGase F glycosidase sensitivity. Immature ER N-glycans are Endo H and PNGase F sensitive; post-Golgi glycans are Endo H resistant. Representative blots and schematic in **c**; Endo H resistant fraction quantified in **d** (n=2, mean ± SEM). **e,** Colocalization of NPC1 variants with the lysosomal marker LAMP1 (Pearson’s coefficient; 4 fields per variant; mean ± SEM). **f,** Representative confocal images of mNG-ALOD4–stained lysosomal cholesterol. Nuclei are in blue. Scale bar: 50 µm. **g,** Cytoplasmic mNG-ALOD4 fluorescence quantified from **f** (200-300 cells per variant, bar shows median, n=3). **h**, Lysosomal membrane integrity assessed by Galectin-3 recruitment to LAMP1-positive structures following LLOMe treatment (8-15 fields per variant, bar shows median). Statistics: (**a,e,g**) one-way ANOVA with Dunnett’s post hoc test versus I1061T; (**h**) one-way ANOVA with Bonferroni’s post hoc test compared to respective untreated control. See also **fig. S5.**

**Figure 4:**
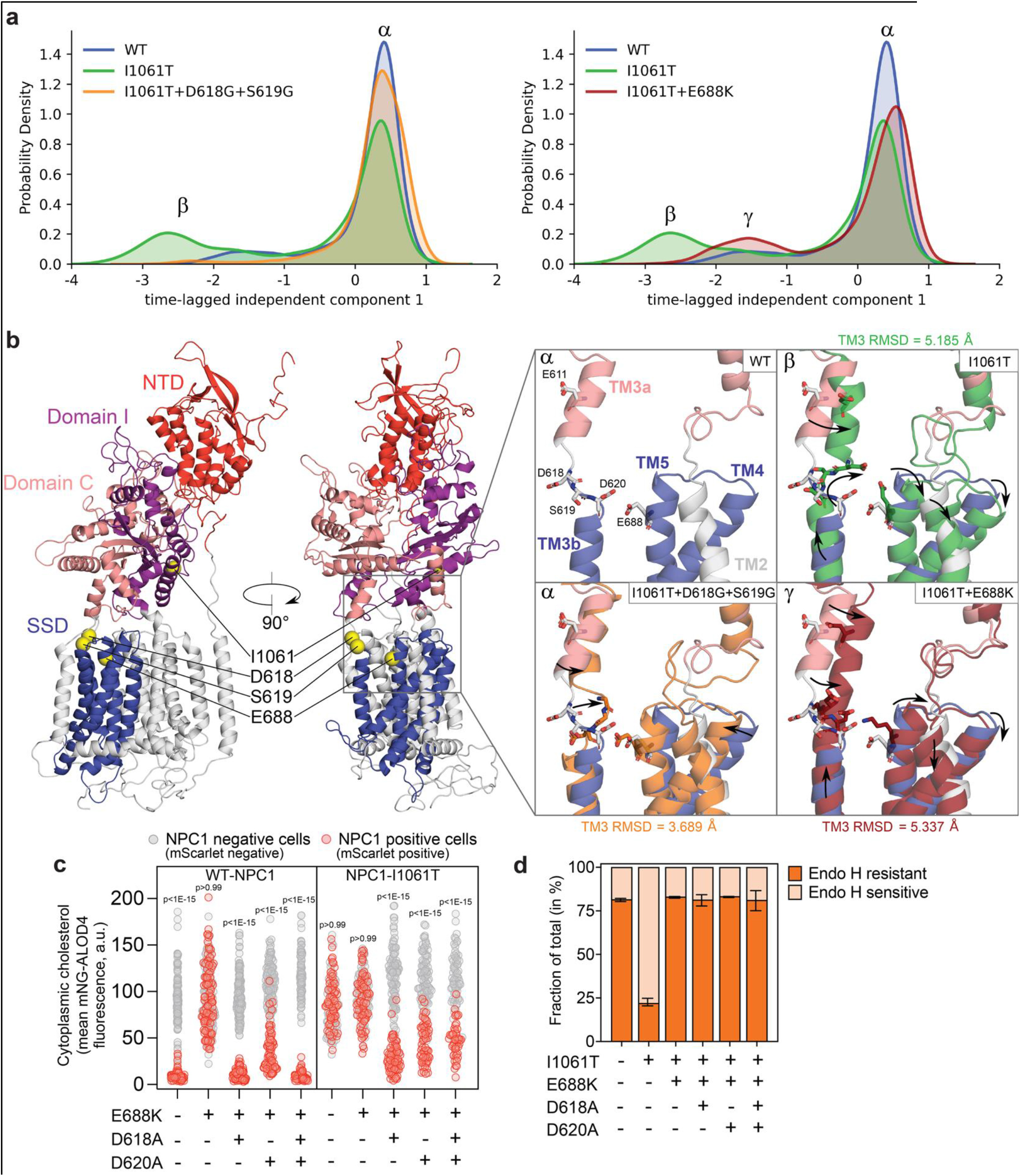
Distinct SSD conformational states underlie folding-only versus functional rescue of NPC1-I1061T by E688K and D618G-S619G. **a**, Time-lagged independent component analysis (tICA) of the SSD from 30 µs all-atom MD simulations of full-length NPC1 embedded in an ER-like lipid bilayer. WT NPC1 predominantly occupies a single conformational state (α, functional). NPC1-I1061T populates an additional aberrant state (β). D618G-S619G restores a WT-like distribution. E688K reduces β occupancy but increases occupancy of an alternate state γ. **b,** Superposition of representative α and aberrant-state conformations from a boxed region centered on the luminal border of the SSD. Arrows indicate displacement of TM helices 3, 4 and 5 in the indicated variants relative to WT. The TM3 RMSD of each depicted variant relative to WT is noted in the panels. **c,** Lysosomal cholesterol (mNG-ALOD4 fluorescence) in *NPC1^-/-^* HeLa cells transiently transfected with constructs encoding NPC1-E688K or NPC1-I1061T-E688K (**fig. S7a** shows the experimental set-up) carrying alanine substitutions at D618, D620, or both. Cytoplasmic mNG-ALOD4 fluorescence was measured in cells expressing NPC1 variants (mScarlet positive, red dots) or untransfected cells (mScarlet negative, gray dots) from the same population (100-200 cells per variant, n=2). **d**, Endo H resistant fraction (a measure of the post-ER fraction) for the variants shown in **c** (mean ± SEM, n=2). Statistics: (**c**) one-way ANOVA with Bonferroni’s post hoc test compared to respective untransfected cells. See also **figs. S6,7** and **Supplementary Table 4**.

The E688K mutation replaces a negatively charged glutamate with a positively charged lysine, introducing the potential for new electrostatic interactions with nearby acidic residues. Consistent with this, the distance between D620 (TM3) and residue 688 (TM5), which are located on opposite sides of the SSD gate, was significantly reduced in the E688K variant compared to all other systems (∼3.5 Å vs. ∼7 Å for WT, **fig. S6a,c**). This electrostatic interaction may stabilize NPC1-I1061T sufficiently to pass ERQC, explaining the rescue of trafficking. However, this interaction may simultaneously lock the SSD in a conformation that prevents the dynamic rearrangements required for cholesterol transport from the tunnel to the SSD or cholesterol binding to the SSD. Structural studies of NCR1 have suggested that the distance between these two residues increases (to ∼7.1 Å) when the gate is open to allow the passage of cholesterol (*30*). Supporting a broader structural impact of this stabilization, E688K was the only corrector mutation that significantly altered the orientation of the N-terminal domain relative to the transmembrane region (**fig. S6b,d**), suggesting that the local constraint propagates to affect global protein architecture. Notably, prior MD studies have suggested that cholesterol binding to the SSD near this region can change NTD dynamics (*38*).

To test whether these local SSD differences propagate to the global communication architecture of NPC1, we constructed residue-residue interaction networks from the simulation trajectories, with nodes representing residues and edges weighted by inter-residue contact frequency (*41–43*). In such networks, edges with high betweenness centrality lie on a disproportionate fraction of inter-residue shortest paths and therefore mark the dominant communication routes through the protein. WT NPC1 and NPC1-I1061T-D618G-S619G shared a continuous high-centrality pathway connecting the NTD to the SSD, whereas this pathway was fragmented in both NPC1-I1061T and NPC1-I1061T-E688K (**fig. S6e**). Network architecture thus tracked with cholesterol-transport function rather than protein abundance or trafficking, supporting the interpretation that E688K stabilizes NPC1-I1061T in a conformation competent for ER exit but incompatible with transport.

To experimentally test the electrostatic interaction predicted by MD simulations, we used a transient transfection assay for NPC1-mediated cholesterol transport in *NPC1*^-/-^ HeLa cells (**fig. S7a**). Expression of WT NPC1 from a weak promoter reduced lysosomal cholesterol only in transfected cells, while NPC1-I1061T failed to do so (**fig. S7b-c**). Each second-site corrector mutation altered lysosomal cholesterol in a manner consistent with our analysis in base-edited eHAP1 cells (compare **Fig.3g** to **fig. S7c**), confirming that these effects reflect the mutations themselves rather than off-target effects of base editing.

Since E688K inactivates both WT NPC1 and NPC1-I1061T, we used both backgrounds to test whether the predicted electrostatic interactions with acidic residues on TM3 were responsible for loss of function. Mutating D618, D620, or both to alanine in either NPC1-E688K or NPC1-I1061T-E688K restored cholesterol transport activity (**Fig.4c,d**). This demonstrates that inactivation by E688K is not caused by loss of the glutamate side chain per se, but rather by a gain of electrostatic interaction between the introduced lysine and neighboring acidic residues, consistent with the compressed D620–K688 distance observed in MD simulations (**fig. S6a**).

### Intrinsic stability of NPC1 variants

To directly measure the intrinsic thermal stability of each NPC1 variant, we performed an in-cell thermal shift assay (CETSA) adapted for membrane proteins using our clonal cell lines expressing each of the NPC1-mScarlet variants(*44*) (**Fig.5a**). In this assay, NPC1-I1061T was markedly less stable than WT NPC1, with a melting temperature (Tm, temperature at which 50% of the protein is lost to aggregation) ∼16 °C lower (**Fig.5b,c**). Reassuringly, reversion of the disease mutation (T1061I) restored thermal stability to WT levels. The four corrector mutations that rescued both trafficking and function showed intermediate thermal stability consistent with a partial restoration of protein folding that is sufficient for ER exit (**Fig.5b,d**). The E688K mutation improved thermal stability to a similar extent as the functional correctors (**Fig.5b,e**), demonstrating that the failure of E688K to rescue cholesterol transport is not related to protein stability. Instead, combined with the MD simulations, these results suggest that E688K stabilizes the protein in a conformation that is compatible with ER exit but incompatible with cholesterol transport. Surprisingly, NPC1-I1061T-Y394C was the least stable variant, with a Tm that was 6 °C lower than even NPC1-I1061T and almost 22 °C lower than WT NPC1 (**Fig.5b,e**). NPC1-I1061T-Y394C remains ER-trapped (**Fig.3c**), likely because it fails ERQC tests due to misfolding. However, its high steady-state abundance suggests that it cannot be recognized and targeted for degradation. This possibility is supported by the fact that NPC1-I1061T-Y394C has a longer half-life than NPC1-I1061T despite remaining in the ER and being less intrinsically stable (**Fig.3b**). Drug profiling showed that NPC1-I1061T-Y394C is an outlier amongst all the NPC1 variants, remaining completely resistant to a panel of drugs that have been shown to improve NPC1-I1061T stability and trafficking (**Fig.5f, fig. S7d**).

**Figure 5:**
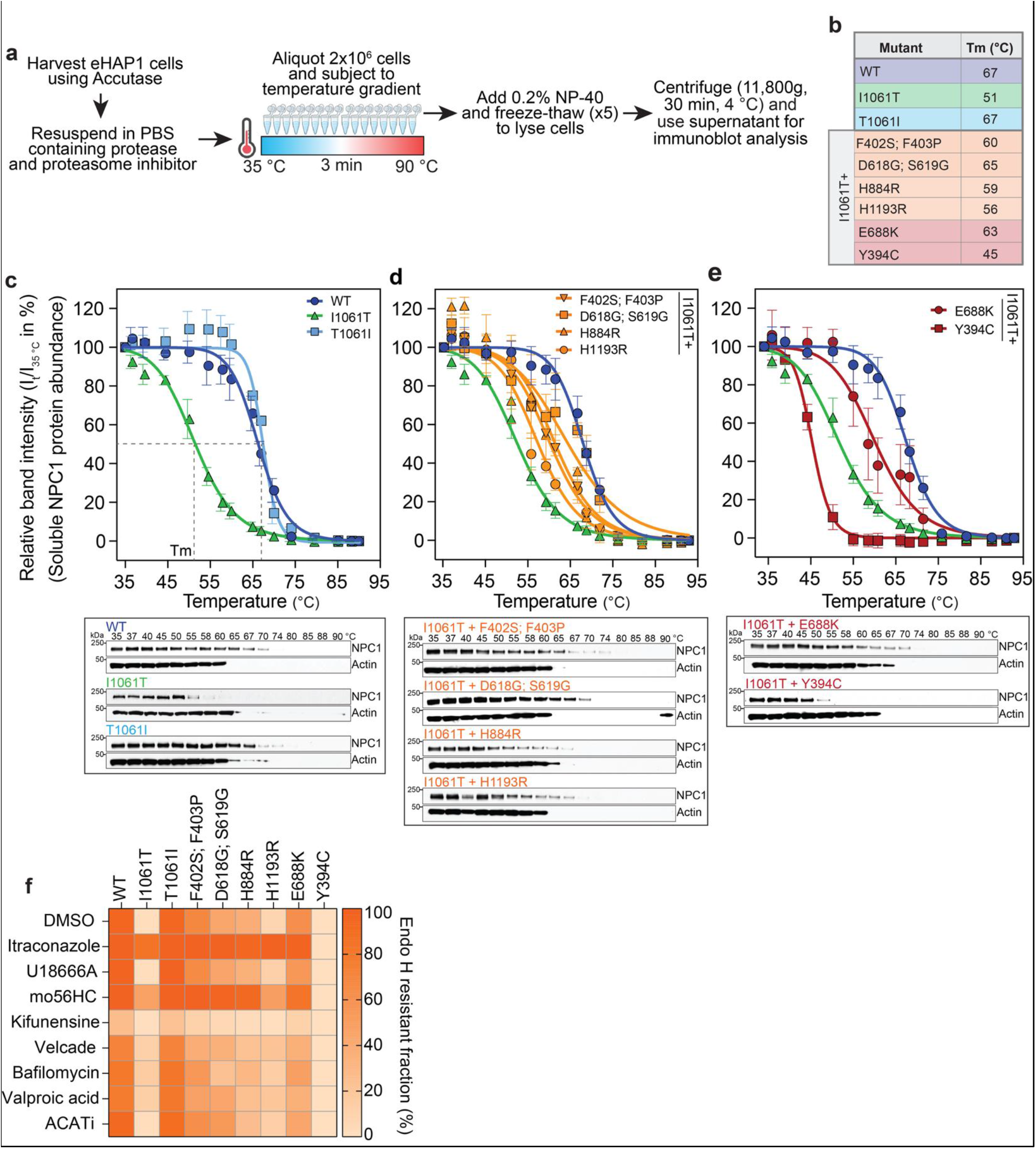
Thermal stability and small molecule responses of NPC1-I1061T corrector variants. **a**, Workflow for the in-cell thermal shift assay (CETSA). **b,** Melting temperatures (Tm, temperature at which 50% of protein is lost to aggregation) for each NPC1 variant. **c-e,** Thermal melting curves (top, mean ± SEM, n=3) and representative immunoblots (bottom) for (**c**) WT, I1061T, and T1061I revertant variant, (**d**) the four correctors that improve both function and abundance on the NPC1-I1061T background, and (**e**) the two correctors that rescue abundance but fail to restore cholesterol transport on the NPC1-I1061T background. Curves plot soluble NPC1 band intensity at each temperature relative to intensity at 35 °C. The uniform melting behavior of Actin across the cell lines serves as a control. **f,** Heatmap of the fraction of each NPC1 variant present in the post-ER compartments (and is thus Endo H resistant) after 24 hr treatment with a panel of pharmacological chaperones known to bind and stabilize NPC1 (1 µM itraconazole, 5 µM U18666A, 1 µM mo56HC), proteostasis modulators (20 µM Kifunensine, 100 nM Velcade, 100 nM Bafilomycin) that inhibit ERAD or ER-phagy pathways, and other agents (4 mM Valproic Acid, 15 µM ACATi) proposed to modulate NPC1 levels. Y394C is resistant to all compounds tested. See also **fig. S7.**

### Corrector mutations rescue NPC1-I1061T in cellular models of Niemann-Pick type C1 disease

To test whether second-site correctors rescue cholesterol transport in disease-relevant cells, we introduced each of the six corrector sgRNAs (**Fig.2c**) into two NPC patient-derived fibroblast lines: one homozygous for I1061T and one compound heterozygote (I1061T/P543L) (**fig. S8**). Because these primary cells were refractory to clonal isolation, we analyzed pooled populations, quantifying both lysosomal cholesterol and editing frequency in parallel (**fig. S8a,b)**. For the four functional corrector mutations, the fraction of cells with reduced lysosomal cholesterol scaled with editing frequency (**Fig.6a**). Rescue was generally lower in the compound heterozygous line, as expected since only the I1061T allele can be corrected. E688K and Y394C failed to reduce cholesterol despite comparable editing, confirming that they stabilize NPC1-I1061T without restoring transport.

**Figure 6:**
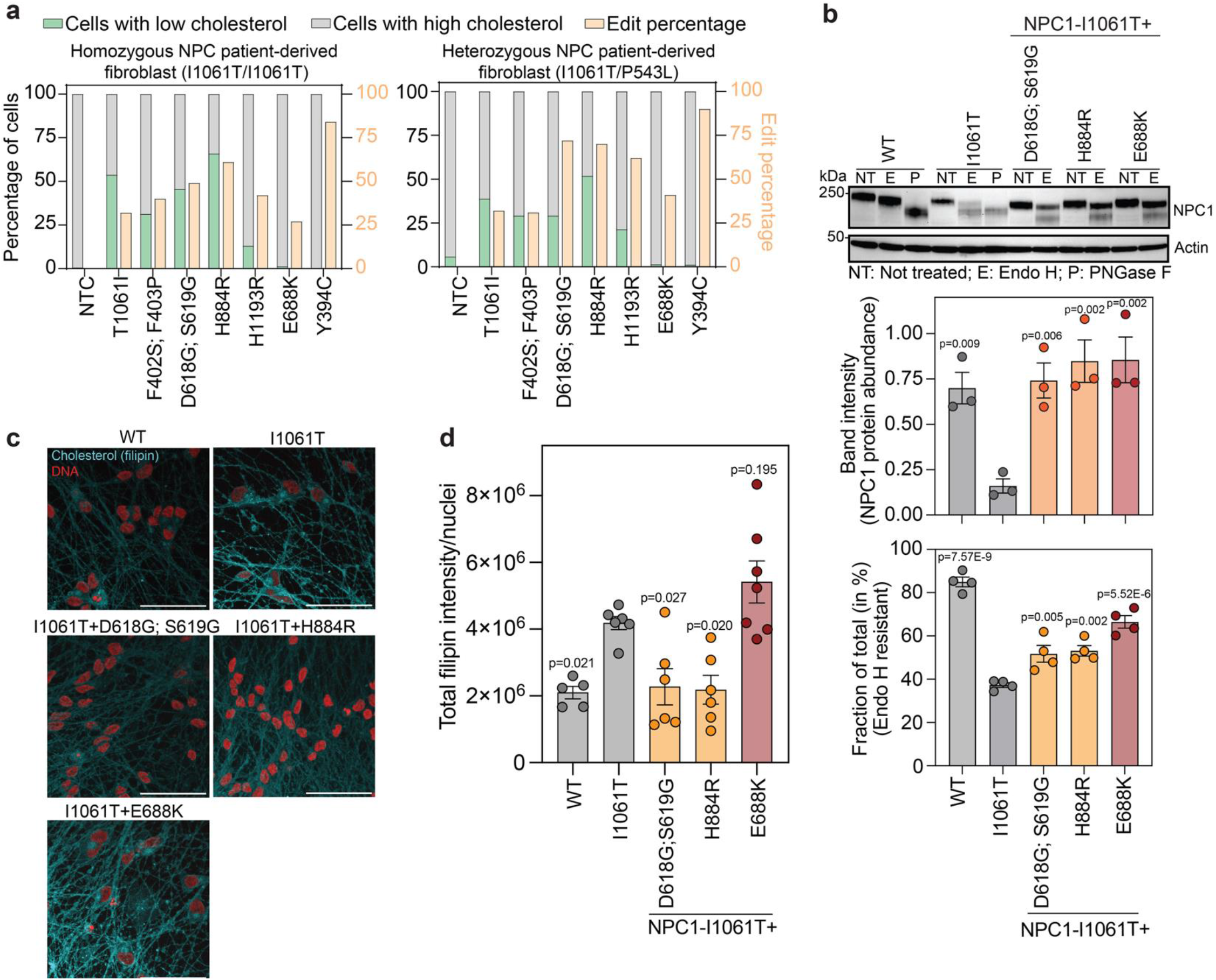
Second-site correctors rescue NPC1-I1061T in NPC patient-derived fibroblasts and iPSC-derived neurons. **a**, Lysosomal cholesterol in two different NPC patient-derived fibroblasts (homozygous I1061T/I1061T and compound heterozygous I1061T/P543L) after installation of corrector mutations by base editing. Lysosomal cholesterol was measured per cell by mNG-ALOD4 staining, and cells were binned as “low cholesterol” (rescued) or “high cholesterol” (unrescued); fractions are plotted on the left y-axis. Base editing percentage for each sgRNA in the population is overlaid for comparison (right y-axis). Data shown is representative of two independent experiments with similar results; see **fig. S8** for amplicon sequencing traces and images showing cholesterol staining. **b,** Immunoblots showing total abundance (NT, Not Treated), Endo H treated (E), and PNGase F treated (P) NPC1-mScarlet variants in lysates made from iNeurons. Actin as a loading control. Bar graphs show quantitation of the Endo H resistant fraction (bottom) (mean ± SEM, n=4) and total abundance (top) (mean ± SEM, n=3). **c,** Filipin staining was used to assess cholesterol abundance in iNeurons expressing the indicated variants of NPC1. Nuclei are in red. Scale bar: 50 µm. **d,** Quantification of filipin fluorescence in iNeuron populations shown in **c** (50-100 cells per variant, mean ± SEM). See **fig. S9a-j** for details of editing and subsequent differentiation of iPSCs into iNeurons, and **fig. S9k,l** for NPC1 abundance and cholesterol accumulation measurements from a second set of iNeurons that were independently differentiated from iPSCs. Statistics: (**b,d**) one-way ANOVA with Dunnett’s post hoc test versus I1061T.

To test correctors in neurons, the cell type whose dysfunction drives NPC disease, we first engineered isogenic induced pluripotent stem cells (iPSCs) with mScarlet-tagged NPC1-I1061T at both endogenous alleles (**fig. S9a-c**). Compared to WT NPC1-mScarlet, NPC1-I1061T-mScarlet was much more Endo H sensitive, showing that it remained trapped in the ER in iPSCs (**fig. S9d,e**). We then introduced three second-site mutations (D618G-S619G, H884R, or E688K) into NPC1-I1061T-mScarlet iPSCs using base editing, isolated clonal cell lines with biallelic edits (**fig. S9f**), and differentiated them into iPSC-derived neurons (iNeurons) using inducible expression of a Neurogenin cassette (*9*) (**fig. S9g,h**). Interestingly, a larger fraction of NPC1-I1061T-mScarlet was Endo H resistant in iNeurons compared to the parental iPSCs, indicating that the ER folding environment may be different between these two cell states (**fig. S9i,j**). Nevertheless, each corrector behaved as expected from our data in eHAP1, HeLa, and patient-derived fibroblast cells. D618G-S619G and H884R increased the total abundance and the Endo H resistant fraction of NPC1-I1061T-mScarlet (**Fig.6b**) and, more importantly, reduced cholesterol accumulation in iNeurons (**Fig.6c,d**; **fig. S9k,l**); E688K increased abundance without reducing cholesterol (**fig. S9k**). In summary, second-site mutations identified in our base editing screens can improve NPC1-I1061T abundance and function across multiple cell types that likely have different capacities for membrane protein folding and quality control in the ER.

### A public corrector mutation that improves the function of structurally diverse misfolded NPC1 variants

To test whether corrector mutations identified in our screens were specific for NPC1-I1061T, we introduced them by base-editing into two additional disease-causing mutations that cause ER retention: NPC1-A1035V and NPC1-R1186H (*45, 46*). A1035V resides in the same luminal domain (domain I) as I1061T, whereas R1186H is located on the opposite side of the membrane in a cytoplasmic loop **(Fig.7a)**. All four corrector mutations partially restored cholesterol transport of NPC1-A1035V, but only D618G-S619G was able to improve the activity of the more severe NPC1-R1186H variant **(fig. S10a,b)**. Second-site correctors therefore differ in their impact, with three acting narrowly across closely related primary mutations and D618G-S619G displaying substantially broader suppressor activity.

**Figure 7:**
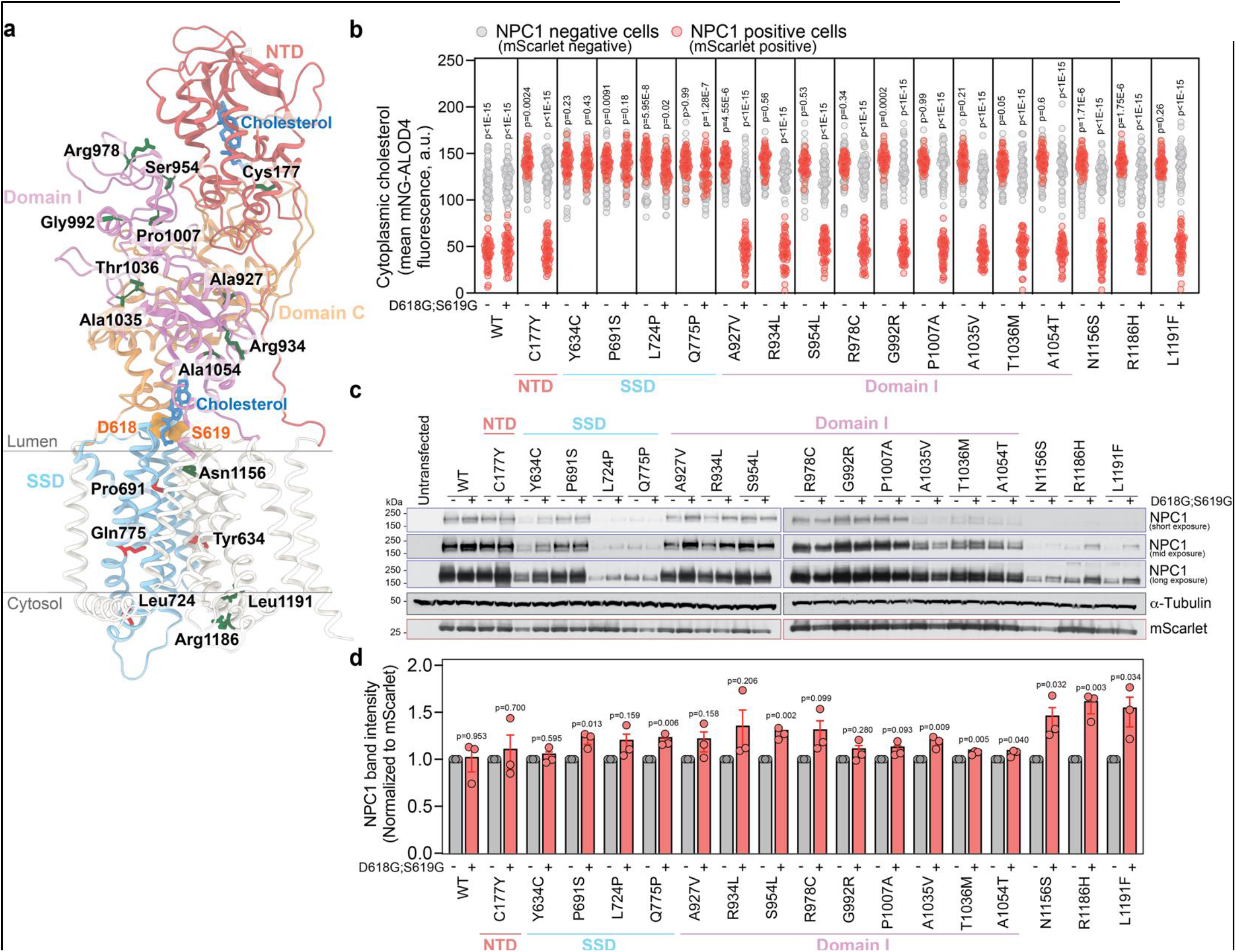
D618G-S619G is a public corrector that rescues structurally diverse pathogenic NPC1 variants. **a**, NPC1 structure (PDB: 6W5U(*29*), bound cholesterol in blue sticks) showing corrector residues D618 and S619 (orange) and 17 missense patient mutations distributed across the NTD (C177Y), SSD (Y634C, P691S, L724P, Q775P), domain I (A927V, R934L, S954L, R978C, G992R, P1007A, A1035V, T1036M, A1054T), transmembrane region (N1156S), and cytoplasmic surface (R1186H, L1191F). Patient mutations are colored according to the rescue of cholesterol levels with corrector mutation (green: rescue, red: no rescue). **b**, Cytoplasmic mNG-ALOD4 fluorescence (lysosomal cholesterol) in *NPC1^-/-^* HeLa cells transiently transfected (see **fig. S7a** for experimental set-up) with NPC1 variants (mScarlet positive, red dots) in the presence or absence of the D618G-S619G corrector mutation in cis. mNG-ALOD4 fluorescence in untransfected cells (mScarlet negative, gray dots) from the same population show high lysosomal cholesterol levels (∼100 cells per variant). **c,d,** Immunoblots and quantification of abundances of the indicated NPC1 protein variants (n=3) in HEK293T cells. Statistics: (**b**) one-way ANOVA with Bonferroni’s post hoc test compared to respective untransfected cells; (**d**) unpaired *t*-test compared to respective control (gray bars). See also **fig. S10.**

To determine whether D618G-S619G functions as a public corrector, we introduced it in cis with 17 additional patient missense mutations spanning the NTD (C177Y), SSD (Y634C, P691S, L724P, Q775P), domain I (A927V, R934L, S954L, R978C, G992R, P1007A, A1035V, T1036M, A1054T), the transmembrane region (N1156S) and cytoplasmic surface (R1186H, L1191F) **(Fig.7a)**. D618G-S619G restored lysosomal cholesterol clearance in 13 of the 17 mutants distributed across all four non-SSD regions tested **(Fig.7b, fig. S10c)**. These included folding-defective alleles (A1035V, R1186H, N1156S (*45, 46*)) as well as mutations spatially remote from D618-S619, such as C177Y ∼62 Å away in the NTD near the cholesterol-binding site and R1186H on the opposite, cytoplasmic face of the membrane. Rescue across such structurally varied positions argues against local, single-domain stabilization and is more consistent with allosteric stabilization of a folding intermediate accessible from multiple primary lesions.

All four rescue failures were SSD mutations (Y634C, P691S, L724P, Q775P), yielding 0/4 at the SSD versus 13/13 rescue elsewhere. P691 sits near the SSD gate and is required for cholesterol transport even in lysosomally-localized NPC1 (*5, 24, 29*). Thus, D618G-S619G functions as a CFTR-style corrector that restores NPC1 folding and trafficking, but likely cannot act as a potentiator to overcome mutations that impair transport activity. Consistent with this view, D618G-S619G increased the steady-state abundance of a majority of the 17 disease mutants of NPC1 **(Fig.7c,d).**

## Discussion

Our base editing screens demonstrate that the folding defect in NPC1-I1061T can be overcome by second-site mutations at diverse positions, establishing a key prerequisite for a therapeutic corrector strategy for Niemann-Pick type C1 disease. The corrector mutations identified here define a variety of structural regions (NTD, domain C, the SSD, and domain I) where binding of a small molecule or a genome edit could, in principle, stabilize the native fold of NPC1-I1061T sufficiently to permit ER exit and restore cholesterol transport. D618G-S619G also rescues 13 additional disease-causing NPC1 mutations suggesting that at least some corrector binding sites may act across multiple patient genotypes, analogous to a “single drug, many alleles” property that has been transformative for CFTR modulators. Our findings nominate the SSD gate region around D618G-S619G (**Fig.2e**) as a candidate single-pocket target capable of addressing a substantial fraction of Niemann-Pick C patient genotypes. Unlike other correctors, D618G-S619G is also in a solvent (and hence drug) accessible region of the protein. The rescue of cholesterol storage defects in patient-derived fibroblasts and induced neurons by corrector mutations provides proof-of-concept for a therapeutic editing or small molecule chaperone strategy.

Our screens revealed that correctors and inactivating mutations converge on residues that encircle a critical bottleneck for cholesterol movement from a protein tunnel to the SSD (*29–31*) (**Fig.2e, inset**). Atomistic simulations and mutagenesis reveal that this region acts as a conformational control point: replacing D618 and S619 with glycines increases the flexibility needed for SSD dynamics and cholesterol transport, while the E688K mutation constrains this same region through an electrostatic interaction with D618 and D620, clamping the SSD in a non-functional state. Opposing mechanisms that either increase or decrease flexibility converge on the same three residues, identifying this site as critical for SSD dynamics. While further work is required to understand how D618G-S619G can rescue diverse mutations across NPC1, our data suggest an allosteric mechanism in which conformational changes initiated at the SSD border propagate through the protein to improve folding efficiency broadly. Consistent with this view, prior MD studies have proposed that cholesterol binding to the SSD (with its hydroxyl positioned near D618, S619 and E688) produces conformational changes that are propagated through the protein to the distant NTD (*38*). One possibility that could explain the global effect of D618G-S619G is that it reduces the probability of kinetically trapped, non-productive folding intermediates, which are recognized and cleared by the ER quality control machinery before they can re-enter the productive folding pathway. A similar mechanism has been proposed for global suppressors of the *Salmonella* Phage P22 tailspike protein (*47*).

Beyond the SSD gate region, a notable general feature of corrector mutations identified in our screens is their proximity to the central tunnel through the center of NPC1 thought to serve as the conduit for cholesterol transport from the NTD to the SSD (**Fig.2a,e**). Interestingly, itraconazole and posaconazole bind inside this tunnel and can act as pharmacological chaperones to improve the folding and lysosomal trafficking of NPC1-I1061T (*11, 32, 36*). Thus, the cholesterol transport tunnel in NPC1 may serve as a structural organizing center for NPC1, a binding region where small molecules or mutations can modulate NPC1 folding, stability, trafficking and function. Of course, such molecules would have to dissociate in the lysosome to allow for cholesterol transport through the tunnel.

An unexpected finding from our screens is that protein stabilization, ER exit, and biochemical function are genetically separable in NPC1. The E688K mutation stabilizes NPC1-I1061T and permits trafficking to the lysosome, but locks the SSD in an aberrant conformation that prevents cholesterol transport. The E688K mutation also inactivates WT NPC1 (**fig. S4**) and thus resembles disease-causing mutations (like P691S) in the SSD that impair NPC1 biochemical activity without impairing its lysosomal trafficking (*5, 24*). Y394C represents a distinct, rarely-reported mechanism: it further destabilizes NPC1-I1061T and causes ER retention yet increases its abundance by evading degradation (by ERAD or ER-phagy). This unexpected separation-of-function mutation shows that ERAD, ER-phagy, and ERQC recognize different properties of a misfolded protein. We note that separation-of-function mutations would not have been identified in a screen based solely on protein abundance or trafficking, highlighting the value of coupling orthogonal phenotypic readouts in suppressor screens. They also provide tools for dissecting how ERQC, ERAD, and ER-phagy distinguish foldable from terminally misfolded membrane proteins, a fundamental question in cell biology that remains poorly understood.

We end by noting that intragenic suppressor screens have a long history in genetics, extending back to work on bacteriophages, and provide unique structural and mechanistic insights into gene and protein function in intact cells (*48, 49*). Our work demonstrates that tiled base editing screens can be used to identify such suppressor mutations in endogenous genes encoding large membrane proteins in mammalian cells. Because every mutation discovered in a base editing screen can be therapeutically installed by the same editor, these screens simultaneously identify corrector mutations and establish the reagents for their deployment, thus providing a general framework for the analysis of other misfolded or conformationally altered disease-causing proteins.

## Materials and Methods

### Reagents and antibodies

Low-density lipoprotein from human plasma (Cat#SAE0053) and filipin (Cat#F4767) were purchased from MilliporeSigma. The following primary antibodies were used for immunoblotting and immunofluorescence: Niemann Pick C1 antibody (Abcam, Cat#ab134113, 1:3000 for immunoblotting, 1:1000 for iPSCs and iNeurons and 1:100 for immunofluorescence), mCherry Polyclonal antibody (Proteintech, Cat#26765-1-AP, 1:500 for immunofluorescence, 1:5000 for immunoblotting), Actin (MilliporeSigma, Cat#MAB1501, 1:10,000 for immunoblotting), beta-Actin Monoclonal Antibody (AC-15) (Invitrogen, Cat#MA191399, 1:2000 for immunoblotting), α-tubulin (Proteintech, Cat#66031-1-Ig, 1:10,000 for immunoblotting), RFP (Rockland, Cat#600-401-379, 1:1000 for immunoblotting), THE™ HA Tag Antibody, mAb, Mouse (GenScript, Cat#A01244-100, 1:1000 for immunoblotting), LAMP1 (Proteintech, Cat#65051-1-Ig, 1:100 for immunofluorescence) and Galectin-3 (Invitrogen, Cat#14-5301-81, 1:100 for immunofluorescence). The following secondary antibodies were used: Peroxidase AffiniPure® Donkey Anti-Mouse IgG (H+L) (Jackson Immunoresearch Laboratories, Cat#715-035-150, 1:10,000), Peroxidase AffiniPure® Donkey Anti-Rabbit IgG (H+L) (Jackson Immunoresearch Laboratories, Cat#711-035-152, 1:10,000), Goat Anti-Mouse IgG (H+L)-HRP Conjugate (Bio-Rad, Cat#1706516, 1:2000), Goat Anti-Rabbit IgG (H+L)-HRP Conjugate (Bio-Rad, Cat#1706515, 1:2000), Donkey anti-Mouse IgG (H+L) Highly Cross-Adsorbed Secondary Antibody, Alexa Fluor™ 488 (Invitrogen, Cat#A-21202, 1:500), Donkey anti-Rat IgG (H+L) Highly Cross-Adsorbed Secondary Antibody, Alexa Fluor™ 594 (Invitrogen, Cat#A-21209, 1:500), and Donkey anti-Rabbit IgG (H+L) Highly Cross-Adsorbed Secondary Antibody, Alexa Fluor™ 594 (Invitrogen, Cat#A-21207, 1:500). Endo H (Cat#P0702L) and PNGase F (Cat#P0708L) were purchased from New England Biolabs. Itraconazole (Sigma-Aldrich, Cat#I6657), U18666A (SelleckChem, Cat#S9669), Kifunensine (Cayman Chemical, Cat#10009437), Bortezomib (Velcade) (LC Laboratories, Cat#B-1408), Bafilomycin A1 (LC Laboratories, Cat#B-1080), Valproic acid (Sigma-Aldrich, Cat#P4543), Sandoz 58-035 (ACATi) (Sigma-Aldrich, Cat#S9318), ROCK inhibitor (Y-27632 dihydrochloride, Cayman Chemical, Cat#10005583).

### Cell culture

eHAP1, a fully haploid human cell line is from Horizon (*20*). The cell line was authenticated by morphology and ploidy analysis as described below to confirm identity. HEK293T was obtained from American Type Culture Collection (ATCC) and its identity was authenticated by the vendor using STR profiling. NPC patient-derived fibroblasts, NPC1^I1061T/I1061T^ (GM18453), and NPC1^I1061T/P543L^ (GM17923) were obtained from the NIGMS Human Genetic Cell Repository at the Coriell Institute for Medical Research. HeLa *NPC1^-/-^* cells were a gift from Suzanne Pfeffer, Stanford University (*52*). All cell lines were confirmed to be mycoplasma-free using an isothermal PCR-based assay (InvivoGen, Cat#rep-mys-10). Experiments were performed using cells between passages 3 and 25. eHAP1 cells were grown in Iscove’s Modified Dulbecco’s Medium (IMDM) containing L-Glutamine (Cytiva, Cat#SH30243FS) and supplemented with 10% fetal bovine serum (FBS) (MilliporeSigma, Cat#F0926), extra glutamine (2 mM) (Cytiva, Cat#SH3003401), and penicillin-streptomycin (50 U/mL) (Gibco, Cat#15140163). HEK293T and human fibroblasts were grown in Dulbecco’s Modified Eagle Medium (DMEM) containing high glucose, with L-glutamine, and sodium pyruvate (Cytiva, Cat#SH30243) and supplemented with 10% FBS, 1x MEM non-essential amino acids solution (Gibco, Cat#11140076), and penicillin-streptomycin (50 U/mL). Cells were subcultured using 0.25% Trypsin-EDTA (Gibco, Cat#25200072) for eHAP1 cells or 0.05% Trypsin-EDTA (Gibco, Cat#25300062) for HEK293T, human fibroblasts and HeLa cells. All cell lines were cultured at 37 °C in a humidified atmosphere containing 5% CO_2_. eHAP1 cells were routinely cultured in 10-Deacetylbaccatin III (Thermo Scientific, Cat#294770010) to enrich for haploid cells and tested for ploidy state by flow cytometry with propidium iodide staining (*53, 54*).

### Recombinant *Sp*Cas9 and sgRNA purification and Ribonucleoprotein (RNP) complex assembly

#### Recombinant SpCas9 Purification

Recombinant *Streptococcus pyogenes* Cas9 (*Sp*Cas9) protein was purified using a protocol adapted from previously described methods (*55*). The expression plasmid pMJ915 (Addgene, plasmid# 69090), which encodes *Sp*Cas9 with an N-terminal His-MBP-TEV tag and a C-terminal 2xSV40NLS sequence, was transformed into Rosetta 2(DE3)pLysS Competent *E. coli* cells (Sigma-Aldrich, Cat#71403). *E. coli* were cultured in Terrific Broth (Thermo Scientific, Cat#H26824.36) at 37 °C. Protein expression was induced with 0.25 mM IPTG at a cell density of OD_600_=0.6, followed by incubation at 18 °C for 16 hr. Cell pellets were harvested by centrifugation and resuspended in lysis buffer (20 mM Tris pH 8.0, 500 mM NaCl, 1 mM TCEP, 5% glycerol, 5 mM imidazole, 0.5 mM PMSF, and 1x SigmaFAST protease inhibitor cocktail (Sigma-Aldrich, Cat#S8820)). Resuspended cells were homogenized using an EmulsiFlex-C5 (Avestin) at 20,000 psi. Lysates were clarified by centrifugation at 50,000g for 30 min, and the supernatant was incubated with His-Tag Purification Resin (Roche, Cat#5893682001) for batch binding. The resin was washed with ten column volumes of wash buffer (lysis buffer excluding PMSF and protease inhibitor cocktail). Bound protein was eluted with an elution buffer (wash buffer supplemented with 250 mM imidazole). TEV protease (homemade) was added to the eluted protein, and the mixture was dialyzed overnight against Ion-Exchange Buffer A (20 mM HEPES pH 7.5, 150 mM KCl, 1 mM TCEP, and 5% glycerol) using a 10 kDa MWCO dialysis cassette (Thermo Scientific, Cat#66453) to cleave the His-MBP tag. The cleaved sample was then applied to a HiTrap SP HP column (Cytiva) equilibrated in 15% Ion-Exchange Buffer B (20 mM HEPES pH 7.5, 1 M KCl, 1 mM TCEP, and 5% glycerol). *Sp*Cas9 was eluted with a linear gradient from 15% to 50% Ion-Exchange Buffer B. Peak fractions were pooled, concentrated using an Amicon ultracentrifugal filter, 30 kDa MWCO (MilliporeSigma, Cat#UFC9030), and subjected to size-exclusion chromatography on a HiLoad 16/60 Superdex 200 prep grade column (Cytiva) equilibrated in Size Exclusion Buffer (20 mM HEPES pH 7.5, 150 mM KCl, 1 mM TCEP, and 10% glycerol). Peak fractions containing purified *Sp*Cas9 were pooled, aliquoted, flash-frozen in liquid nitrogen, and stored at −80 °C.

#### sgRNA Synthesis and RNP Assembly

sgRNA template assembly, amplification, *in vitro* transcription, and purification were performed as described previously (*55*). *In vitro* transcription was carried out using the HiScribe T7 High Yield RNA Synthesis Kit (New England Biolabs, Cat#E2040S) following the manufacturer’s instructions. Transcription reactions were subsequently treated with RNase-free DNase I (Thermo Scientific, Cat#EN0521) to remove DNA template, followed by Quick CIP (New England Biolabs, Cat#M0525). sgRNAs were purified using the RNeasy Mini kit (Qiagen, Cat#74104). For CRISPR/Cas9-mediated HDR experiments, Cas9-RNP complexes were assembled immediately before use.

### Endogenous tagging of *NPC1*

To generate an eHAP1 clonal cell line with an endogenously tagged *NPC1* allele, *Sp*Cas9 protein was complexed with sgRNA (5′-CAGTCAGGATGCCCTGCGAG-3′) and electroporated with the HDR template plasmid (pUC19-NPC1/mScarlet with ∼800-bp left and right homology arms flanking mScarlet for C-terminal tagging of NPC1, see **Supplementary Table 5**). For electroporation, Cas9–RNP complex was prepared by mixing 100 pmol of *Sp*Cas9 protein with 120 pmol of sgRNA in Cas9 buffer (20 mM HEPES pH 7.4, 150 mM KCl, 1 mM MgCl_2_, 1 mM TCEP, and 10% glycerol) and 1 μg of the HDR template plasmid for 10 min. The RNP complex was added to 500,000 low-passage eHAP1 cells and electroporated using a NEPA21 Electro-Kinetic transfection system (Nepagene). The specific electroporation parameters used were: poring pulse of 190 V, poring length of 5 ms, and the remaining parameters were set as default. The cells were incubated in antibiotic-free medium with 1 μM of HDR Enhancer v2 (Integrated DNA Technologies, Cat#10007910). 24 hr after electroporation, growth medium containing 1 μM of HDR Enhancer v2 was added. Approximately 72 hr post-electroporation, single cells were sorted into 96-well plates, gating for cells with top 0.1% mScarlet fluorescence intensity using a SH800S Cell Sorter (Sony). Single cell clones were subsequently expanded in culture. Genotyping was performed using multiple primer pairs complementary to the mScarlet sequence, homology arms and the genomic region outside the homology arms to confirm the correct site of integration in the *NPC1* gene locus (**Supplementary Table 5)**. Expression of the NPC1-mScarlet fusion protein was confirmed by immunoblotting using antibodies against mScarlet and NPC1.

### Precise genome editing of *NPC1* locus in eHAP1 cells

sgRNAs and single-stranded oligonucleotide (ssODN) donor templates for precise genome editing of the *NPC1*locus via HDR were designed using the CRISPR HDR Design Tool (Integrated DNA Technologies). The targeting sequences for NPC1 were 5’-GACATTACTGGCTATAAGTC-3’ (I1061T), 5’-GTGGTAGGTCATGAAGTACG-3’ (A1035V) and 5’-CCTCTTCCGCGCGCTCCACG-3’ (R1186H). ssODN donor templates were designed to introduce either the I1061T, A1035V or the R1186H mutations into the *NPC1* gene. The sequences of the ssODN templates are listed in **Supplementary Table 5**. A silent mutation was incorporated into the ssODN for NPC1-I1061T to introduce a HpaI restriction site to facilitate screening, designed using silent mutagenesis tool (molbiotools). eHAP1 cells or eHAP1 NPC1-mScarlet cells were subjected to electroporation to deliver the CRISPR/Cas9 RNP complex and ssODN donor templates for HDR. Electroporation was performed as described in the *Endogenous tagging of NPC1* section using the NEPA21 Electro-Kinetic transfection system. Single-cell sorting was performed approximately 72 hr post-electroporation to isolate individual cells for clonal expansion. Sorting was performed based on viability gating. Clones were screened and confirmed for the presence of the desired NPC1 mutations by genotyping. Briefly, genomic DNA was extracted, and the region of the target NPC1 locus was amplified by PCR. The PCR product was then digested with restriction enzyme (New England Biolabs, see **Supplementary Table 5**) to confirm the intended HDR event. Putative positive clones were confirmed by Sanger sequencing of the targeted NPC1 locus.

### Purification of recombinant His-mNeonGreen-FLAG-ALOD4 (mNG-ALOD4)

The recombinant cholesterol-binding protein domain was purified according to a protocol adapted from (*23*). The expression plasmid pRSET B_His6-mNeonGreen-FLAG-ALOD4 was transformed into Rosetta 2(DE3)pLysS competent *E. coli* cells (Sigma-Aldrich, Cat#71403). Cells were cultured in Lysogeny Broth (LB) containing 10 g/L Tryptone, 5 g/L Yeast Extract, and 10 g/L NaCl (pH 7.0) supplemented with 100 µg/mL carbenicillin at 37 °C. Upon reaching an OD_600_ = 0.6, the temperature was reduced to 18 °C, and protein expression was induced with 1 mM IPTG for 18 hr. Cells were harvested by centrifugation at 3,500 g for 15 min at 4 °C. The cell pellets were resuspended in Buffer A (50 mM Tris-HCl pH 7.5, 150 mM NaCl, and 1 mM DTT) supplemented with 1 mM PMSF and 1x SigmaFAST protease inhibitor cocktail (Sigma-Aldrich, Cat#S8820). The suspension was homogenized using an EmulsiFlex homogenizer (Avestin) via two passes at 15,000 psi. The lysate was clarified by centrifugation at 25,000g for 30 min at 4 °C, and the resulting supernatant was incubated with 3 mL of IMAC Resin (Roche, Cat#5893682001) on an orbital shaker at 4 °C for 30 min. The resin-lysate mixture was loaded into a gravity-flow column. The column was washed with 50 mL of Buffer A, followed by a second wash with 50 mL of Buffer A containing 5 mM imidazole. Bound protein was eluted using Buffer B (Buffer A supplemented with 250 mM imidazole, pH 7.5). Peak fractions were pooled and concentrated to a volume of 3 mL using an Amicon Ultra-15 Centrifugal Filter Unit (10 kDa NMWL) (MilliporeSigma, Cat#UFC901024D). To remove protein aggregates, the concentrated eluate was subjected to centrifugation at 20,000g for 30 min at 4 °C. The sample was then injected onto a Superdex 200 column equilibrated in Buffer A using an ÄKTA pure FPLC system (Cytiva). Peak fractions were resolved on 4-12% NuPAGE™ Bis-Tris protein gel (Invitrogen, Cat#WG1402BOX) and stained with GelCode Blue Stain Reagent (Thermo Scientific, Cat#24592). Final pooled fractions were supplemented with 20% glycerol, and the protein concentration was determined using Pierce BCA protein assay (Thermo Scientific, Cat#23225). Purified protein aliquots were flash-frozen in liquid nitrogen and stored at −80 °C.

### Cholesterol staining using mNG-ALOD4 for flow cytometry

eHAP1 mutant library cells were subjected to a fluorescence-activated cell sorting (FACS) assay using the accessible cholesterol sensor ALOD4. Cells were seeded and grown to 80% confluency. Growth media was replaced with complete IMDM supplemented with 25 µg/mL LDL and incubated for 16 hr. Following the LDL pulse, cells were treated with 5 mM hydroxypropyl-β-cyclodextrin (HPCD, Thermo Scientific, Cat#297560250) for 15 min at 37 °C to minimize plasma membrane cholesterol levels. Cells were harvested by trypsinization and collected by centrifugation. The cells were fixed with 4% paraformaldehyde (PFA) for 10 min on ice, followed by quenching with 0.125 M glycine for 5 min on ice. Cells were permeabilized by 0.6% Triton X-100 for 10 min on ice, followed by two washes in PBS. Permeabilized cells were resuspended in staining buffer containing 1 µM mNG-ALOD4, 1% BSA in PBS and incubated on a slow rocker for 1 hr at 4 °C. Following incubation, cells were pelleted and resuspended in PBS containing 1 mM EDTA for flow cytometry.

### FACS-based CRISPR base editing screens

#### Library construction

For sgRNA library design, the Beagle tool from the Broad Institute Genetic Perturbation Platform (https://portals.broadinstitute.org/gppx/beagle/public) was used to design guides for the PAM-flexible (NG-PAM) adenine and cytosine base editors. The sgRNA library tiles each exon across *NPC1* such that each codon is targeted by multiple sgRNAs. The design strategy also allowed sgRNAs to extend ∼20 nucleotides into introns at each intron-exon junction to create splice-site mutations. The resulting library was filtered to exclude guide sequences containing BsmBI restriction sites. The Ensembl transcript ID used for guide design was ENST00000269228.10. BsmBI recognition sites and appropriate overhangs were appended to each gRNA sequence for Golden Gate cloning into base-editor plasmids (pRDA_429, Addgene, plasmid# 179098 expressing ABE8e Cas9-NG and pRDA_336, Addgene, plasmid# 179095 expressing BE3.9 Cas9-NG). Specific primer sites were utilized for the selective amplification of the subset of NPC1 targeting sequence from a larger, multiplexed oligonucleotide synthesis pool. The final oligonucleotide sequence was as follows 5’-[Forward Primer]*CGTCTCACACCG*[targeting sequence, 20 nt]*GTTTCGAGACG*[Reverse Primer]-3’. Oligonucleotide pools were synthesized by GenScript using the GenTitan™ High-Fidelity 92k platform. PCR amplification of the oligonucleotide pool was performed using 25 μL of 2x NEBNext PCR Master Mix (New England Biolabs, Cat#M0541S), 2 μL of the oligonucleotide pool (∼2 ng), 5 μL of primer mix at a final concentration of 0.5 μM, and 18 μL of nuclease-free water. PCR cycling conditions were: (1) Initial denaturation: 98 °C for 1 min; (2) 98 °C for 10 s; (3) 62 °C for 10 s; (4) 72 °C for 20 s; (5) repeat steps 2-4 for 18 cycles; (6) Final extension: 72 °C for 2 min. The following primer (5’-3’) pairs were used for amplification: NPC1-ABE: Forward: AGGCACTTGCTCGTACGACG, Reverse: ATGTGGGCCCGGCACCTTAA NPC1-CBE: Forward: GTGTAACCCGTAGGGCACCT, Reverse: GTCGAGAGCAGTCCTTCGAC Control: Forward: TTACTAGACTGCGGGGCCAC, Reverse: CGGCTCGATCTGCGACGAAT The resulting amplicons were resolved on a 2% agarose gel, purified by Zymoclean Gel DNA Recovery Kit (Zymo Research, Cat#D4008) and ligated into the pre-digested base-editor plasmids using BsmBI-v2 (New England Biolabs, Cat#R0739L) and T4 DNA ligase (New England Biolabs, Cat#M0202S) following the Golden Gate cloning method. The ligation product was isopropanol precipitated and electroporated into Stbl4 electrocompetent cells (Invitrogen, Cat#11635018) and grown at 30 °C. Plasmid DNA was prepared (HiSpeed Plasmid Maxi Kit, Qiagen, Cat#12662) and guide representation and distribution in the libraries were confirmed by next-generation sequencing (NGS).

#### Mutant cell library preparation

Lentivirus for pooled library production was generated by transfecting HEK293T cells (20×10^6^) in a 225 cm^2^ flask with a DNA mixture containing packaging plasmids psPAX2 (50 μg; Addgene, plasmid# 12260), pMD2.G (5 μg; Addgene, plasmid# 12259), and 40 μg of ABE or CBE plasmid library, using polyethylenimine (PEI, Polysciences, Inc., Cat#23966-2) at 1:3 ratio. The supernatant containing lentivirus was collected 48 hr post-transfection and filtered (0.45 μm). To determine lentiviral titer, eHAP1 cells were transduced with varying volumes of lentivirus. 48 hr post-infection, puromycin (Sigma-Aldrich, Cat#P8833) was added at a concentration of 5 μg/mL. After five days, cell viability was assessed to determine the viral dose that resulted in a transduction efficiency of 20-30%, corresponding to a multiplicity of infection (MOI) of less than 0.35. To generate mutant library cells, eHAP1 NPC1-I1061T or eHAP1 NPC1-I1061T-mScarlet cells were transduced in two biological replicates at a low MOI of 0.3. Briefly, 1.5×10^6^ cells were seeded per well in a 12-well plate containing IMDM. 24 hr post seeding, plates were subjected to spinfection with lentivirus mixed with polybrene (8 μg/mL) at 640g for 2 hr to enhance transduction efficiency. Cells were incubated at 37 °C for 6 hr, after which they were trypsinized, pooled and transferred into 225-cm^2^ flasks each for ABE and CBE libraries. Two days post-spinfection, puromycin (5 μg/mL) was added and maintained for seven days to eliminate non-transduced cells. After selection, the cell libraries were frozen and stored in liquid nitrogen.

#### FACS-based phenotype enrichment for pooled screens

For each biological replicate screen, mutant library cells were thawed in 15-cm dishes, split and seeded for the FACS sorting stage. For the function screen, eHAP1 NPC1-I1061T cells were prepared according to section *Cholesterol staining using mNG-ALOD4 for flow cytometry* and then sorted using Sony SH800S cell sorter to isolate the bottom 10% of the cell population with low mNG-ALOD4 fluorescence. Collection tubes (15 mL) were pre-coated with 1 mL FBS for at least 1 hr at 4 °C. Sorted cells were collected in 5 mL PBS supplemented with 2% FBS. For the abundance screen, eHAP1 NPC1-I1061T-mScarlet cells were harvested using 0.25% Trypsin-EDTA, pelleted at 500g for 5 min and resuspended in IMDM. Live cells were subsequently sorted into PBS containing 2% FBS to isolate the top 10% of the cell population with high mScarlet fluorescence. For each replicate, 4–6 million cells were collected per screen, corresponding to ∼2,350-3,500× representation of the ∼1,700 guide library (including 278 non-targeting controls). An unsorted population from each replicate was retained as a reference for log-fold change (LFC) normalization. After sorting, cells were pelleted (4,800g, 10 min, 4 °C) and stored at −80 °C prior to genomic DNA extraction.

#### Next Generation Sequencing (NGS)

Genomic DNA (gDNA) was isolated from sorted and unsorted cell populations using the Mag-Bind Blood & Tissue DNA HDQ Kit (Omega Bio-Tek, Cat#M6393). For PCR amplification of the integrated sgRNA sequences, 1-2 μg of gDNA was added to a 50 μL reaction mixture containing NEBNext High-Fidelity 2X PCR Master Mix (New England Biolabs, Cat#M0541L) and primers containing adaptor sequences with unique barcoded i7 and i5 indexes for Illumina sequencing (See **Supplementary Table 5**). PCR cycling conditions were: (1) Initial denaturation: 95 °C for 1 min; (2) 94 °C for 30 s; (3) 52.5 °C for 30 s; (4) 72 °C for 30 s; (5) repeat steps 2-4 for 25 cycles; (6) Final extension: 72 °C for 5 min. PCR products were resolved on a 2% agarose gel, purified by Zymoclean Gel DNA Recovery Kit and concentrations were estimated using Qubit 1X dsDNA HS Assay Kit (Cat#Q33231). The samples were then sequenced on an Illumina MiSeq instrument using the MiSeq reagent kit v3 (150 cycles, single end, 25M reads) using a custom sequencing primer (5’-CCGATCTGCTTTATATATCTTGTGGAAAGGACGAAACACCG-3’).

#### Data analysis

Data processing and analysis were performed in R (v4.3.2). Raw sequencing reads were demultiplexed and converted to FASTQ files using the bcl2fastq module (v2.20.0), with each file assigned to its corresponding condition based on the 8-nt P5 and P7 barcoded primers. Guide RNA sequences (20 nt) were extracted from FASTQ files using the PreprocessReads function from the QuasR package (v1.36.0) (*56*). Reads were aligned to a custom reference containing all NPC1 and non-targeting control library guide sequences using the Subread algorithm (Rsubread v2.16.2) (*57*). Aligned reads were summarized into a count matrix, which was normalized to reads per million (RPM) for each condition: RPM = (reads per sgRNA/total reads per condition)x10^6^. To prevent issues with zero counts, a log_2_ transformation was applied to the RPM values after adding a pseudocount of 1 to all counts. Guides with log₂-normalized RPM in the unsorted population outside 1.5x of the interquartile range (IQR) were filtered out to remove outliers and poorly represented sgRNAs. Log-fold change (LFC) between sorted and unsorted populations was computed for each sgRNA. To standardize enrichment of sgRNAs in the sorted population, z-scores were calculated using LFC distributions from non-targeting control guides (*25*). One-sided *p*-values for enriched sgRNAs were derived from the upper tail of the standard normal distribution, and FDR-adjusted p-values were obtained using the Benjamini-Hochberg method. As a second independent method, we also used the well-established MAGeCK pipeline for analysis of screen results (*27, 28*). Using custom R functions, predicted base editing outcomes (N3-N10 window) were classified into mutation categories (nonsense, splice-site, missense, silent) based on library annotations. These categories were incorporated into the dataset for downstream analysis and visualization. Z-scores were calculated for each replicate, and replicate reproducibility was assessed *via* z-score correlation plots. All scripts and analysis pipelines are available on GitHub, ensuring transparency and reproducibility of the results.

### Validation of base-editor sgRNA hits in eHAP1 cells

To validate the top hits from the base editing screens, sgRNAs with z-scores ≥2 were selected. Individual targeting sequences were cloned into either the pRDA_429 or pRDA_336 base-editor plasmid. Lentivirus for each cloned sgRNA was generated by transfecting HEK293T cells (1.5×10^6^ in 6-cm dish) with a DNA mixture containing packaging plasmids psPAX2 (4 μg; Addgene, plasmid# 12260), pMD2.G (0.8 μg; Addgene, plasmid# 12259), and 4 μg of sgRNA plasmid, using polyethylenimine (PEI, Polysciences, Inc., Cat#23966-2) at 1:3 ratio. The supernatant containing lentivirus was collected 48 hr post-transfection and filtered (0.45 μm). For each sgRNA, 1.5×10^6^ eHAP1 NPC1-I1061T-mScarlet cells were seeded per well in a 12-well plate. 24 hr post cell seeding, cells were transduced with 1.5 mL of lentivirus in a growth medium containing polybrene (8 μg/mL). The plates were centrifuged at 640g for 2 hr before transferring to a 37 °C incubator. After 6 hr, cells were trypsinized and transferred to a 10-cm dish. Puromycin selection (5 μg/mL) was initiated 48 hr post-transduction and continued for 5-7 days. Following selection, cells were plated and assessed for rescue of NPC1 function and abundance. Cholesterol clearance (NPC1 function) was assessed by flow cytometry as described in the section *Cholesterol staining using mNG-ALOD4 for flow cytometry*. NPC1 protein abundance was determined by measuring mScarlet fluorescence intensity by flow cytometry on a Sony SH800S cell sorter.

### FACS-based isolation of clonal mutant lines

For the sgRNAs selected after validation, the allele frequency in the base-edited eHAP1 NPC1-I1061T-mScarlet cell population was estimated by NGS. gDNA was extracted from population corresponding to each individual sgRNA using QuickExtract DNA Extraction Solution (Lucigen, Cat#QE09050), and the target loci were amplified using a two-step PCR protocol. The initial PCR utilized site-specific primers appended with Illumina adapters, followed by a second PCR to incorporate unique barcodes for demultiplexing (see **Supplementary Table 5**). FASTQ files from NGS were processed using CRISPResso2(*58*) to quantify the allele frequencies and verify the predicted base-edits. To generate clonal lines for detailed characterization, the base-edited mixed populations were single-cell sorted into individual wells of a 96-well plate. Single cell clones were subsequently expanded in culture. Following expansion, the genotype of each resulting clone was validated by Sanger sequencing of the targeted locus to confirm the specific mutation.

### Immunoblotting

All cells were harvested by scraping. Cells were lysed in RIPA Lysis Buffer (50 mM Tris pH 8.0, 150 mM NaCl, 1% NP-40, 0.1% SDS, 0.1% Deoxycholate, 10% glycerol and 0.5 mM dithiothreitol (DTT)) containing 1x SigmaFAST protease inhibitor cocktail (Sigma-Aldrich, Cat#S8820). Samples were vortexed for 30 min at 4 °C to ensure complete lysis. Lysates were then clarified by centrifugation at 20,000g for 30 min at 4 °C, and the resulting supernatant was collected for analysis. Protein concentrations were determined using Pierce BCA Protein Assay Kit (Thermo Scientific, Cat#23225) with BSA standards. For immunoblotting, 4x NuPAGE™ LDS Sample Buffer (Invitrogen, Cat#NP0007) containing 50 mM DTT was added to 30 μg of total protein, and incubated for 30 min at 37 °C. Proteins were resolved on 4-12% NuPAGE™ Bis-Tris Protein Gels (Invitrogen, Cat#WG1402BOX) using 1x MOPS-SDS running buffer (50 mM MOPS, 50 mM Tris Base, 0.1% SDS, 1 mM EDTA, pH 7.7) and transferred to nitrocellulose membrane (Bio-Rad, Cat#1620112). Membranes were blocked for 30 min at room temperature in Blocking Buffer (BB) (Tris-buffered saline (TBS) containing 0.1% Tween-20 (TBS-T) and 5% non-fat dry milk). Membranes were incubated with primary antibodies diluted in BB, overnight at 4 °C on a slow rocker. After three washes with TBS-T, membranes were incubated for 1 hr at room temperature with the appropriate HRP-conjugated secondary antibody diluted in BB. After three washes with TBS-T, chemiluminescence was detected using Pierce ECL Western Blotting Substrate (Thermo Scientific, Cat#32209) or SuperSignal West Pico PLUS Chemiluminescent Substrate (Thermo Scientific, Cat#34580), and captured on film.

### Endoglycosidase H (Endo H) assay

The Endo H assay was modified from (*59*). Briefly, three reactions (negative control, Endo H, PNGase F) were set up per sample containing 20 µg cell lysate (30 µg for I1061T), 1 µl glycoprotein denaturing buffer, and nuclease-free water to make up to 10 µl volume and incubated at 37 °C for 15 min. After incubation, the Endo H samples received 2 µl 10x G3 reaction buffer, 5 µl water, and 3 µl Endo H to make a total reaction volume of 20 µl. In the negative control reactions, the Endo H was replaced with water. PNGase F reactions received 2 µl of 10x G2 reaction buffer, 2 µl 10% NP-40, 3 µl water, and 3 µl PNGase F. All reactions were placed at 37 °C for 3 hr. Reactions were terminated with 7 µl 4x NuPAGE™ LDS Sample Buffer containing 50 mM DTT and incubated for 10 min at 37 °C. Proteins were resolved on 4-12% NuPAGE™ Bis-Tris Protein Gels followed by immunoblot as described above.

### Cycloheximide assay

Cells were treated with 60 μg/mL cycloheximide, harvested at 0, 4, 8 and 12 hr timepoints and processed according to the section *Immunoblotting*.

### Fluorescence imaging and analysis

#### mNG-ALOD4 staining and cholesterol analysis

Cells grown on coverglass were pulsed with 20 µg/mL LDL in growth medium for 16 hr. Following the LDL pulse, cells were washed with PBS and fixed with 4% PFA for 10 min on ice. Residual fixative was quenched with 0.125 M glycine for 10 min on ice. Fixed cells were permeabilized with 0.1% Triton X-100 for 10 min on ice. For cholesterol staining, cells were incubated with 1 µM mNG-ALOD4 in 1% BSA for 1 hr on ice. Nuclei were counterstained with DAPI (ProLong Diamond Antifade Mountant, Invitrogen, Cat#P36962) during mounting. Images were acquired on a Leica TCS SP8 confocal microscope equipped with a 63× oil-immersion objective. Cytoplasmic fluorescence intensities of the mNG-ALOD4 channel were quantified using CellProfiler (*60*) (v 4.2.8). Nuclear masks were first defined by automated thresholding of the DAPI channel using the minimum cross-entropy method. Cytoplasmic compartments were then identified by radial expansion of the nuclear boundaries while excluding neighboring nuclei. To avoid signal contribution from the plasma membrane, the resulting cytoplasmic masks were further shrunk by 4 pixels. Mean fluorescence intensities were extracted from these cytoplasmic compartments for quantification.

#### Filipin staining and cholesterol analysis

Cells grown on coverglass were pulsed with 20 µg/mL LDL in growth medium for 16 hr. Following the LDL pulse, cells were washed with PBS and fixed with 4% PFA for 20 min at room temperature. To quench residual PFA, cells were incubated with 0.125 M glycine for 10 min at room temperature. Free cholesterol was subsequently stained using 0.05 µg/mL filipin + 5% FBS in PBS for 2 hr at room temperature. Cells were washed three times with PBS and mounted with NucSpot® 640 mounting media (Biotium, Cat#23016), used as a nuclear counterstain. Mid-plane section images of cells were acquired using a Leica TCS SP8 confocal microscope. To minimize photobleaching of the UV-sensitive filipin signal, laser power and scan speeds were optimized. Cytoplasmic fluorescence intensities were quantified using CellProfiler (*60*) (v 4.2.8). Nuclear masks were defined by automated thresholding of the NucSpot® 640 channel using the minimum cross-entropy method. Cytoplasmic masks were subsequently generated by radial expansion of the nuclear boundaries while excluding neighboring nuclei. To avoid signal contribution from the plasma membrane, the resulting cytoplasmic masks were further shrunk by 4 pixels. Mean fluorescence intensities were extracted from these cytoplasmic compartments for quantification.

#### Lysosomal localization of NPC1

Cells grown on coverglass were washed with PBS and fixed with ice-cold 100% methanol for 20 min at −20 °C. Following three PBS washes, cells were incubated in a saponin-based blocking buffer (SBB) (0.02% saponin, 5% normal donkey serum, and 1% BSA in PBS) for 30 min at room temperature. Primary antibodies against LAMP1 (1:100) and NPC1 (1:100) were diluted in SBB and incubated for 1 hr at room temperature. Following three washes with PBS containing 0.02% saponin, cells were incubated with species-specific secondary antibodies (Donkey anti-Mouse Alexa Fluor 488 (1:500, for LAMP1) and Donkey anti-Rabbit Alexa Fluor 594 (1:500, for NPC1)) diluted in SBB for 1 hr at room temperature. Cells were washed with PBS containing 0.02% saponin and mounted using ProLong Diamond Antifade Mountant with DAPI. Images were acquired on a Leica TCS SP8 confocal microscope. To capture the full spatial distribution of NPC1 and LAMP1, z-stacks encompassing the entire cell volume were collected at 0.5 µm intervals. Colocalization analysis was performed on the resulting image stacks using the Just Another Colocalization Plugin (JACoP) within ImageJ/Fiji (*61*).

#### Lysosomal damage assay and Galectin-3 recruitment

To assess lysosomal membrane integrity, cells were treated with DMSO (vehicle) or indicated doses of LLOMe for 1 hr to induce lysosomal rupture. Following treatment, cells were washed three times with PBS and fixed with ice-cold 100% methanol for 20 min at −20 °C. Cells were permeabilized and blocked with SBB for 1 hr on ice. For dual-labeling of damaged lysosomes, cells were incubated for 2 hr on ice with primary antibodies against Galectin-3 (1:100) and LAMP1 (1:100) diluted in SBB. Following three washes in PBS containing 0.02% saponin, cells were incubated with species-specific secondary antibodies (Donkey anti-Mouse Alexa Fluor 488 (1:500, for LAMP1) and Donkey anti-Rat Alexa Fluor 594 (1:500, for Galectin-3)) diluted in SBB for 1 hr on ice. Cells were washed with PBS containing 0.02% saponin and mounted using ProLong Diamond Antifade Mountant with DAPI. Confocal z-stacks (0.5 µm intervals) were acquired on a Leica TCS SP8 confocal microscope. To quantify the extent of lysosomal damage, Manders’ overlap coefficients were calculated from the fraction of LAMP1 signal coinciding with Galectin-3 puncta using the Just Another Colocalization Plugin (JACoP) (*61*) in ImageJ/Fiji.

### In Cell Thermal Shift Assay (CETSA)

Cells were harvested with Accutase cell detachment solution (MilliporeSigma, Cat#SCR005) and collected by centrifugation at 340g for 5 min. Cells were washed twice with PBS and resuspended in PBS supplemented with 1x SigmaFAST protease inhibitor cocktail, 20 mM N-ethylmaleimide (NEM) and 1 μM bortezomib (LC Laboratories, Cat#B-1408). 20 µl of cell suspension containing 2×10^6^ cells was distributed into a series of 0.2 mL PCR tubes and the aliquots were subjected to a 3 min heat shock across a thermal gradient ranging from 35-90 °C, followed by rapid cooling to 25 °C in a thermal cycler. To facilitate protein extraction, 5 µl of 1% NP-40 (final concentration 0.2%) was added to each tube. Cells were lysed *via* five consecutive freeze-thaw cycles, consisting of snap-freezing in liquid nitrogen for 3 min followed by thawing in a 25 °C water bath for 3 min. Insoluble proteins and cellular debris were removed by centrifugation at 11,800g for 30 min at 4 °C. The resulting supernatants containing the soluble protein fraction were resolved on 4-12% NuPAGE™ Bis-Tris Protein Gels and immunoblotted according to the section *Immunoblotting*.

### cDNA-mediated expression of NPC1 variants in HeLa NPC1*^-/-^* cells

To evaluate the functional capacity of NPC1 variants, HeLa *NPC1^-/-^* cells (*52*) were transiently transfected with constructs encoding the indicated NPC1 variants and a P2A-linked mScarlet reporter under the control of a truncated EF1α core promoter (212 bp). WT NPC1 cDNA was obtained from Addgene, plasmid# 164972. Constructs containing various mutations were generated using site-directed mutagenesis using primers specified in **Supplementary Table 5**. Transfections were performed using Lipofectamine 3000 (Invitrogen, Cat#L3000008) according to the manufacturer’s instructions. Cholesterol distribution was evaluated by mNG-ALOD4 staining followed by confocal imaging (as described in the section *Fluorescence imaging and analysis*: *mNG-ALOD4 staining and cholesterol analysis*). Presence of P2A self-cleaving peptide between NPC1 and mScarlet enabled identification of transfected cells by mScarlet fluorescence. To assess the rescue of NPC1 variants in HeLa *NPC1^-/-^* cells, the population was first classified as mScarlet-positive (minimum intensity ≥30%; expressing the NPC1 variant) or mScarlet-negative (maximum intensity <30%; non-expressing) based on mScarlet fluorescence intensity within the cytoplasmic mask. The mean mNG-ALOD4 fluorescence intensity was then extracted from the cytoplasmic regions of both populations for plotting.

### Immunoblotting of NPC1 variants in HEK293T cells

The same constructs described above (NPC1 variants with a C-terminal 3xHA tag followed by a P2A-linked mScarlet reporter) were used to assess NPC1 expression by immunoblot. HEK293T cells were transiently transfected using Lipofectamine 3000 according to the manufacturer’s instructions. Whole-cell lysates were resolved on 4-12% NuPAGE™ Bis-Tris protein gels and immunoblotted as described in the section *Immunoblotting*. NPC1 was detected using THE™ HA Tag Antibody, and cleaved mScarlet was detected using an anti-mCherry polyclonal antibody.

### Validation in NPC1*^I1061T/I1061T^* and NPC1*^I1061T/P543L^*human primary fibroblasts

For each sgRNA, 50,000 fibroblasts were seeded per well in a 12-well plate. 24 hr post cell seeding, cells were transduced with 1.5 mL of lentivirus in a growth medium containing polybrene (8 μg/mL). The plates were centrifuged at 640g for 2 hr before transferring to a 37 °C incubator. After 6 hr, cells were trypsinized and transferred to a 6-well plate. Puromycin selection (2 μg/mL) was initiated 48 hr post-transduction and continued for 5-7 days. Following selection, cells were assessed for editing efficiency and rescue of cholesterol accumulation. gDNA was isolated using QuickExtract DNA Extraction Solution (Lucigen, Cat#QE09050) according to the manufacturer’s instructions. The targeted mutation locus was amplified by PCR using the primers specified in **Supplementary Table 5** and analyzed by Sanger sequencing. Editing efficiency and the percentage of edited alleles were quantified using EditR (https://moriaritylab.shinyapps.io/editr_v10/). The fibroblasts were stained with mNG-ALOD4 (as described in the section *Fluorescence imaging and analysis*: *mNG-ALOD4 staining and cholesterol analysis*) and imaged to quantify the percentage of cells with high and low cholesterol. The minimum intensity observed in NTC fibroblasts (∼0.14 in 0-1 scale) was defined as the threshold separating low- and high-cholesterol populations. For each guide, the percentage of cells exhibiting low cholesterol staining was calculated as the fraction of cells with mean mNG-ALOD4 intensity below this NTC-derived threshold.

### Generation, characterization, and analysis of NPC1-mScarlet-I reporter iPSC and iNeuron models

#### Generation and characterization of NPC1-mScarlet-I reporter iPSC lines

Homology directed repair was used to insert an mScarlet-I tag after the final *NPC1* exon. 2E iPSCs (from human foreskin fibroblasts) were transfected with an *NPC1*-targeting donor repair template containing mScarlet-I flanked by 916 bp 5′ and 1077 bp 3′ homology arms and 2 Cas9/sgRNA plasmids with EGFP marker targeting the terminal NPC1 exon/3′ UTR region (See **Supplementary Table 5**). Homology-directed repair resulted in insertion of the mScarlet-I coding sequence immediately downstream of the final NPC1 exon. Genotyping primers positioned outside the homology arms were used to distinguish wild-type and tagged alleles. Positively transfected cells were selected via FACS and plated at single cell density. High quality clonal colonies (2 homozygous for mScarlet-I, 1 homozygous WT) were selected and genotyped. Clones were sequenced at the *NPC1* locus to confirm faithful insertion. Pluripotency and CNV were assessed for the selected clones.

#### NGN2 and I1061T insertion and validation of doxycycline-inducible NGN2 NPC1 iNeuron models

TALENs were used to insert human *NGN2* into the CLYBL safe harbor locus of the homozygous mScarlet-I iPSCs. An mNeonGreen marker was used to manually select single cells. Cell clones were selected and genotyped to confirm insertion. To ensure successful integration, colonies were subjected to doxycycline induction and expression of neuronal markers was confirmed. CNV was assessed to ensure cell line integrity. CRISPR/Cas9 was then used to introduce the I1061T mutation into the iPSCs following our previous methods and confirmed by sequencing (*59*). The resulting isogenic iPSC lines contained doxycycline-inducible NGN2 expression from CLYBL and mScarlet-tagged NPC1 protein carrying either the wild-type allele (WT-mSc) or the I1061T mutation (I1061T-mSc). For quantitative reverse transcription polymerase chain reaction (qRT-PCR) analysis RNA was collected using TRIzol (Ambion, Cat#15596018); cDNA was synthesized using cDNA High Capacity Reverse Transcription Kit (Applied Biosystems, Cat#4368814); and PCR using TaqMan probes (Thermo Fisher Scientific, for 18S (4310893), SYN1 (Hs01018066), MAP2 (Hs00258900), and MAPT (Hs00902194)) in ABI 7900HT Sequence Detection System. Immunofluorescence staining for MAP2 (primary antibody against MAP2, 1:150, MilliporeSigma Cat#MAB3418/clone AP20; secondary: Alexa Fluor 488 goat anti-mouse IgG (H+L), 1:500, Invitrogen, Cat#A11029) confirmed robust neuronal differentiation and neurite outgrowth. MAP2-positive cells were quantified using ImageJ/Fiji, and the percentage of MAP2-positive cells is reported.

#### Base editing of NPC1-I1061T-mScarlet-I reporter iPSC lines

I1061T-mSc iPSCs were cultured to approximately 70% confluency, dissociated using Accutase, and resuspended in DPBS (-/-). For each electroporation reaction, 10^6^ cells were resuspended in 100 µL of Buffer R (Invitrogen, Cat#N10096) containing 1 µg of an editing plasmid expressing a blue fluorescent protein (BFP) marker (pRDA_429, Addgene, plasmid# 179098 expressing ABE8e Cas9-NG and pRDA_336, Addgene, plasmid# 179095 expressing BE3.9 Cas9-NG, in both plasmids puromycin gene was replaced with BFP) and electroporated using the Neon NxT Transfection System. Electroporation conditions consisted of three pulses at 1200 V for 10 ms. Following electroporation, cells were plated in pre-warmed StemFlex medium (Gibco, Cat#A33493-01) supplemented with ROCK inhibitor (1:2000). Media was replaced after 2 hr, and cells were left undisturbed for 24 hr. The following day, conditioned media was collected, and cells were dissociated using Gentle Cell Dissociation Reagent. BFP-positive cells were single-cell sorted by FACS into Geltrex-coated (Gibco, Cat#A4000046803) 96-well plates containing a 1:1 mixture of conditioned media and fresh StemFlex medium supplemented with ROCK inhibitor (1:2000). iPSCs were maintained in 96-well plates with daily media changes until colonies reached confluency. A small aliquot of cells from each clone was retained for Sanger sequencing to assess the presence of the desired edits.

#### iNeuron differentiation

iNeurons were generated from iPSCs following previously established protocols (*59*). Briefly, iPSCs were plated onto Geltrex-coated dishes in StemFlex medium supplemented with ROCK inhibitor on day −1. The following day (day 0), cells were treated with StemFlex medium containing doxycycline (1:2000) to induce neuronal differentiation. On day 2, neural progenitors were dissociated using Accutase and replated onto fresh Geltrex-coated dishes in 3N medium supplemented with doxycycline (1:2000). Media was changed daily on days 3 and 4 by replacing half of the conditioned media with an equal volume of fresh 3N medium containing doxycycline (*59*). On day 5, media was removed, and cells were collected in DPBS (-/-) followed by cell lysis. Endoglycosidase H (Endo H) and PNGase F digestions were performed as previously described (*59*) to assess NPC1 glycosylation status and determine the relative abundance of mature and immature glycoforms.

#### Cell lysis and protein quantification

Cells were harvested by scraping and lysed in RIPA buffer (Teknova, Cat#R3792) with 10 mM NEM (n-ethylmaleimide) (Sigma-Aldrich, Cat#E3875-5G) and cOmplete Mini tablet (MilliporeSigma, Cat#11836153001). Cells were then sonicated in RIPA lysis buffer for 2-4 cycles. Following sonication, the lysates were centrifuged at 12,000g for 8 min at 4 °C to pellet DNA. The supernatants were collected, and the protein concentration was determined with the DC^TM^ Protein Assay Kit (Bio-Rad, Cat#5000111) with BSA standards.

#### Endo H assay

Three reactions including negative control, Endo H digestion, and PNGase F digestion were prepared per sample. Cell lysates with 16 µg of total protein were prepared per reaction, 3 µl of 10x glycoprotein denaturing buffer (New England Biolabs, Cat#B1704S) was added per reaction, and nuclease-free water to make a final volume of 30 µl. The reactions were then incubated at 37 °C for 15 min. Following incubation, in the negative control reactions, 4 µl of 10x glycobuffer 3 (New England Biolabs, Cat#B1720S), and nuclease-free water were added to reach a final volume of 40 µl. Endo H reactions received 4 µl of 10x glycobuffer 3, 2 µl of Endo H, and nuclease-free water to reach a final volume of 40 µl. Finally, the PNGase reactions received 4 µl of 10x glycobuffer 2 (New England Biolabs, Cat#B3704S), 4 µl of 10% NP-40 (New England Biolabs, Cat#B2704S), 1.5 µl of PNGase F, and water to reach a final volume of 40 µl. All reactions were incubated at 37 °C for 3 hr. Following incubation, to terminate the reaction, 16 µl of 4x NuPAGE^TM^ LDS Sample Buffer containing 167 mM of DTT was added, and immunoblotting was performed.

#### Immunoblotting

Proteins were resolved on 4-12% NuPAGE^TM^ Bis-Tris Midi Gels (Invitrogen, Cat#WG1401BX10). The blots were run for 2 hr at 130 V using 1x MOPS-SDS running buffer (Thermo Scientific, Cat#J62847.K2) and transferred using the semi-dry transfer method at 13 V for 2 hr to an Immobilon-P 0.45 µM PVDF membrane (Millipore, Cat#IPVH85R). Membranes were blocked for 5 min in Everyblot Blocking Buffer (Bio-Rad, Cat#12010020) at room temperature with gentle rocking. Membranes were then incubated in primary antibodies diluted in Everyblot Blocking Buffer for 1 hr at room temperature on a slow rocker. The membranes were then washed five times for 5 min each in TBS-T. Following the washes, the membranes were then incubated for 1 hr at room temperature with the appropriate HRP-conjugated secondary antibody diluted in Everyblot Blocking Buffer. After five washes for 5 min each in TBS-T, chemiluminescence was detected using Pierce ECL Western Blotting Substrate (Thermo Scientific, Cat#32106).

#### Filipin staining of iNeurons

Cells grown on coverglass were washed with PBS and fixed with 4% PFA for 20 min at room temperature. Free cholesterol was subsequently stained using 0.05 µg/mL filipin + 5% FBS in PBS for 2 hr at room temperature. Cells were washed three times with PBS and mounted with NucSpot® 640 mounting media (Biotium, Cat#23016), used as a nuclear counterstain. Z-stacks were acquired using a Leica TCS SP8 confocal microscope. Maximum intensity projections were generated from each z-stack. Raw integrated density was quantified from the projected images using ImageJ/Fiji and divided by the number of nuclei for each frame to obtain normalized filipin fluorescence.

#### NPC1 staining in iNeurons

Cells grown on coverglass were washed with PBS and fixed with 4% PFA for 20 min at room temperature, permeabilized and blocked (blocking buffer: 1×PBS, 1% normal donkey serum, 10 mg/mL BSA, 0.1% Triton X-100) for 30 min. Cells were incubated with rabbit anti-mCherry primary antibody (to detect NPC1-mScarlet-I) in blocking buffer for 2 hr at room temperature, washed three times in PBS+0.1% Triton X-100, and incubated with Alexa Fluor 594-conjugated donkey anti-rabbit secondary antibody (1 hr, room temperature). After washing three times with PBS+0.1% Triton X-100, coverslips were mounted with NucSpot® 640 mounting media. Z-stacks were acquired using a Leica TCS SP8 confocal microscope.

### Molecular Dynamics (MD) Simulations

#### Simulation Setup

Since NPC1 folds in the ER, simulations were carried out at neutral pH using an ER-like symmetric membrane. As a starting structure for simulations we considered two prior NPC1 structures solved at neutral pH: PDB ID: 6UOX (*32*) (4.02 Å) and PDB ID: 6W5S (*29*) (3.0 Å); the RMSD between these two for 1067 pruned atom pairs is 0.896 angstroms. We chose to use 6UOX because of superior wwPDB metrics, including clashscore, Ramachandran outliers, and sidechain outliers. The ligand (Itraconazole) in 6UOX was removed because none of our experiments contained Itraconazole and, more importantly, Itraconazole acts as a pharmacological chaperone itself (*11*) and stabilizes NPC1-I1061T; hence, its inclusion in the simulations would potentially confound the effects of our second-site suppressors. We also did not include any cholesterol ligands because of the low cholesterol content of the ER (compared to the plasma membrane or lysosome) (*37, 62*).

All unresolved residues in the full protein sequence were modeled using MODELLER (*63*) in accordance with UniProt entry O15118-1. All generated residues exist either at the beginning of the NTD or within the cytoplasmic domain between transmembrane helices and were not constrained to any specific secondary structure. MODELLER was similarly used to generate all mutant systems (**Supplementary Table 4**). To determine protonation states for acidic residues under physiological conditions, the pKa was calculated using PROPKA 3 for each residue (*64, 65*). Residues with a pKa greater than physiological pH 7.2 were protonated accordingly (**Supplementary Table 4**). Residues were glycosylated based on commonly resolved glycans in select fully resolved NPC1 structures (**Supplementary Table 4**). The structures examined include 3JD8, 6UOX, 6W5R, 6W5S, 6W5T, 6W5U and 6W5V. The proteins were embedded in a membrane bilayer and solvated using CHARMM-GUI (*66, 67*) with lipid content corresponding to a mammalian endoplasmic reticulum membrane (*37*) (**Supplementary Table 4**). The systems were solvated with TIP3P water (*68*) and 150 mM KCl to mimic physiological conditions, with a water gap of 25 Å and box width of 125 Å to prevent self-interaction. Hydrogen masses were repartitioned to 3.024 Da using ParmEd’s HMassRepartition (*69*) to enable simulations with a 4 fs timestep (*70*). Atomic interactions were characterized using the CHARMM36m forcefield (*71*). The total number of atoms in each system is shown in **Supplementary Table 4**.

#### Pre-production simulations

All pre-production simulations were performed using the OpenMM 7.6 biomolecular simulation package (*72*). Each mutant was prepared with 10 replicates. Systems were minimized with OpenMM’s LocalEnergyMinimizer to a tolerance of 5.0 kJ/mol, which uses the limited-memory Broyden-Fletcher-Goldfarb-Shanno algorithm (L-BFGS)(*73*). Systems were then heated from 0 to 310 K over 10 ns under NVT conditions using the Langevin thermostat (*74*). The protein backbone atoms were constrained using a force constant of 10 kcal mol^-1^ Å^-2^, and the phosphorous atoms of the phospholipid head groups were constrained using a force constant of 0.5 kcal mol^-1^ Å^-2^ to ensure membrane stability. Systems were then equilibrated under NPT conditions for 10 ns at 310 K and 1 atm using OpenMM’s MonteCarloMembraneBarostat with harmonic restraints only on the protein backbone, followed by 10 ns relaxing said constraints. Lastly, the systems were equilibrated in the same thermodynamic ensemble for 10 ns without any restraints.

#### Production Simulations

Production simulations were run from equilibrated frames from each system. Systems were all simulated using periodic boundary conditions. The Particle Mesh Ewald method (PME) (*75*) was used for computing long-range electrostatic interactions with a cutoff of 12 Å for non-bonded interactions. Each system was run with 30 clones per replicate. Each trajectory was run for 25,000,000 steps at 4 fs per step with a save rate of 25,000 steps, yielding 100 ns trajectories with 0.1 ns per frame. Across 300 total trajectories per system, this yielded 30 µs of simulation data per mutant for a total of 240 µs simulated.

#### Trajectory analysis

Simulation trajectories were processed and stripped of water molecules using cpptraj(*76*) as part of the AmberTools25 software. Analysis scripts were written in Python using MDTraj(*77*) for computing pairwise residue distances/domain angles. Python packages NumPy (*78*), SciPy (*79*), pandas (*80*), seaborn (*81*), and matplotlib (*82*) were used for general scientific computations and plot visualizations. Deeptime (*83*) was used to implement time-lagged Independent Component Analysis (tICA) (*40, 84, 85*), the details of which are discussed below. NetworkX (*86*) was used to generate residue interaction networks (RIN) using in-house scripts based on MD-TASK (*43*), the details of which are also discussed below. Protein structure and network visualizations were generated using PyMOL (The PyMOL Molecular Graphics System, Version 3.0 Schrödinger, LLC.). For each trajectory, the NTD angle or SSD distance was averaged over frames 201-1000 (first 20 ns discarded as equilibration), and these per-trajectory averages were plotted across mutants.

#### Dimensionality Reduction using tICA

time-lagged Independent Component Analysis (tICA) was used to process MD data into a more interpretable form by reducing the trajectories into linear combinations of calculated collective variables. Input features were calculated as pairwise distances between residues in the SSD, generated using the following protocol for a total of 16,290 residue-pair distances, of which 1,629 were retained for analysis:

resi_pairs_full = []

\# SSD defined as residue indices 618-798

res_start, res_end = 618, 798

for i in range(res_start, res_end+1):

for j in range(i+1, res_end+1):

resi_pairs_full.append((i,j))

\# retain every 10th pair due to memory constraints

resi_pairs_tica = resi_pairs_full[::10]

A tICA lagtime of 20 ns was chosen, corresponding to implied timescale calculations indicating system stability at said lagtime.

#### Network Construction and Analysis

A residue interaction network (RIN) was built for each system, with nodes representing amino acid residues and edges representing contact frequency between residues. A network was generated for each frame of each trajectory, and residues were considered in contact if their β-carbons (α-carbons for glycine) were within 6.7 Å. These networks were then concatenated across each frame of each trajectory for a given system, yielding one representative network for each mutant with every node corresponding to a residue, and every edge and its weight corresponding to the contact and contact frequency between two residues across the ensemble of simulations. To reduce noise in the representative networks, weak edges with a weight of less than 0.01 were dropped from the networks. Additionally, to make the networks more representative of the dynamic contacts and conformational changes in the system (*42*), edges with a weight of 0.95 or greater were also dropped from the networks, allowing for the networks to better represent the flow of information through the protein rather than the static contacts that do not contribute to overall dynamics. Betweenness centrality metrics were calculated using NetworkX’s edge_betweenness_centrality function to determine the importance of a given edge in protein conformation. Major and minor pathways were determined based on edge betweenness centrality, with major pathways involving the highest centrality edges in a continuous sequence, and minor pathways branching off from the major pathway included if their edge betweenness centrality was at least 20% of the maximum edge betweenness centrality for a given network.

## Data visualization, statistics and reproducibility

Quantitative data were visualized using GraphPad Prism (v11.0.0). Structural figures were generated using UCSF ChimeraX (v 1.9). PDB codes for structural figures are provided in the respective figure legends. Immunoblot panels and microscopy images were processed using ImageJ/Fiji (v1.54p). Flow cytometry data were analyzed using Floreada (https://floreada.io/). FASTQ files from NGS were processed using CRISPResso2 (*58*). Sanger sequencing traces were visualized and quantified using SnapGene (v8.2.2) and EditR (https://moriaritylab.shinyapps.io/editr_v10/). Figure panels were assembled using Adobe Illustrator (v30.2.1). All n values cited in the legends refer to biological replicates. Statistical significance was determined using one-way ANOVA followed by pairwise comparisons between groups using Dunnett’s post hoc test versus one control group or Bonferroni’s post hoc test when compared to their respective control group. For two sample comparisons, significance was assessed by unpaired Student’s *t*-tests. Specific *p*-values are reported directly in the figure panels.

## Data availability

All raw and processed sequencing data supporting the CRISPR-based screening and subsequent analyses are available in the Sequence Read Archive (SRA) repository under BioProject accession number PRJNA1429439. The specific read counts and processed data files derived from the initial base editing screen are listed in **Supplementary Tables 1 and 2**. All source data and corresponding raw quantification files used for the validation of sgRNAs are provided in **Supplementary Table 3**, accompanying the paper.

Primary and processed data for the atomistic MD simulation is available for reviewers from the following link: https://databank.illinois.edu/datasets/IDB-3753443?code=Lhbg4qp6q8WoM_OfTUwqJTbrNAydh0eD9q6FHilG56w. This data package will be made public at the time of publication.

## Code availability

The custom R scripts and computational pipelines utilized for the processing and statistical analysis of the base editing screen data were written in R (v4.3.2) and are publicly available at the following GitHub repository: https://github.com/RohatgiLab/Base_editing_analysis_tool.

## Competing interests

The authors declare no competing interests

## Author contributions

Conception: S.R., P.S., G.V.P., R.R. Design of study: S.R., P.S., G.V.P., R.R., M.L.S., E.G., D.S. Data collection: S.R., P.S., E.G., G.V.P., E.P., M.S., T.S., R.A. Data analysis: S.R., P.S., G.V.P., M.P., E.G., M.L.S. Supervision: R.R., M.L.S., D.S. Acquisition of funding: R.R., M.L.S., D.S. Writing and editing: S.R., P.S., R.R. with input from all authors.

## Acknowledgements

This work was supported by grants from the Ara Parseghian Medical Research Fund at the University of Notre Dame (to R.R.) and the NIH (R35GM142745 to D.S.; R35GM154959 to M.L.S. and T32MH019113-31 to M.S.). We also gratefully acknowledge Dr. Brian Blagg from the Warren Center for Drug Discovery at University of Notre Dame.

## Supplementary Materials

figs. S1 to S10

Tables S1-S5

## Supplementary Tables

**Supplementary Table 1.** Comprehensive presentation of function screen. Screen data were analyzed using a *z*-score method based on a control distribution of ∼280 non-targeting sgRNAs. An alternative analysis using MAGeCK is also provided(*27, 28*). Note that sgRNAs were selected for validation and mechanistic studies (as in **Fig.2b**) using the z-score method, not MAGeCK.

**Supplementary Table 2.** Comprehensive presentation of abundance screen. Screen data were analyzed using a *z*-score method based on a control distribution of ∼280 non-targeting sgRNAs. An alternative analysis using MAGeCK is also provided(*27, 28*). Note that sgRNAs were selected for validation and mechanistic studies (as in **Fig.2b**) using the z-score method, not MAGeCK.

**Supplementary Table 3.** Screen results were validated (as outlined in **fig. S3a**) by individually introducing sgRNAs enriched in the FACS gates used for the screens (**Fig.1d**) into eHAP1 NPC1-I1061T-mScarlet cells. This table provides all the data from these validation studies.

**Supplementary Table 4.** Summary of molecular dynamics simulation parameters, including simulated systems and atom counts, protonated residues, glycosylation sites and glycans, and membrane lipid composition.

**Supplementary Table 5.** Oligonucleotides used in this study, including Sanger sequencing primers, NGS primers, demultiplexing barcodes, HDR templates, sgRNAs and cloning primers.

---

**fig S1:**
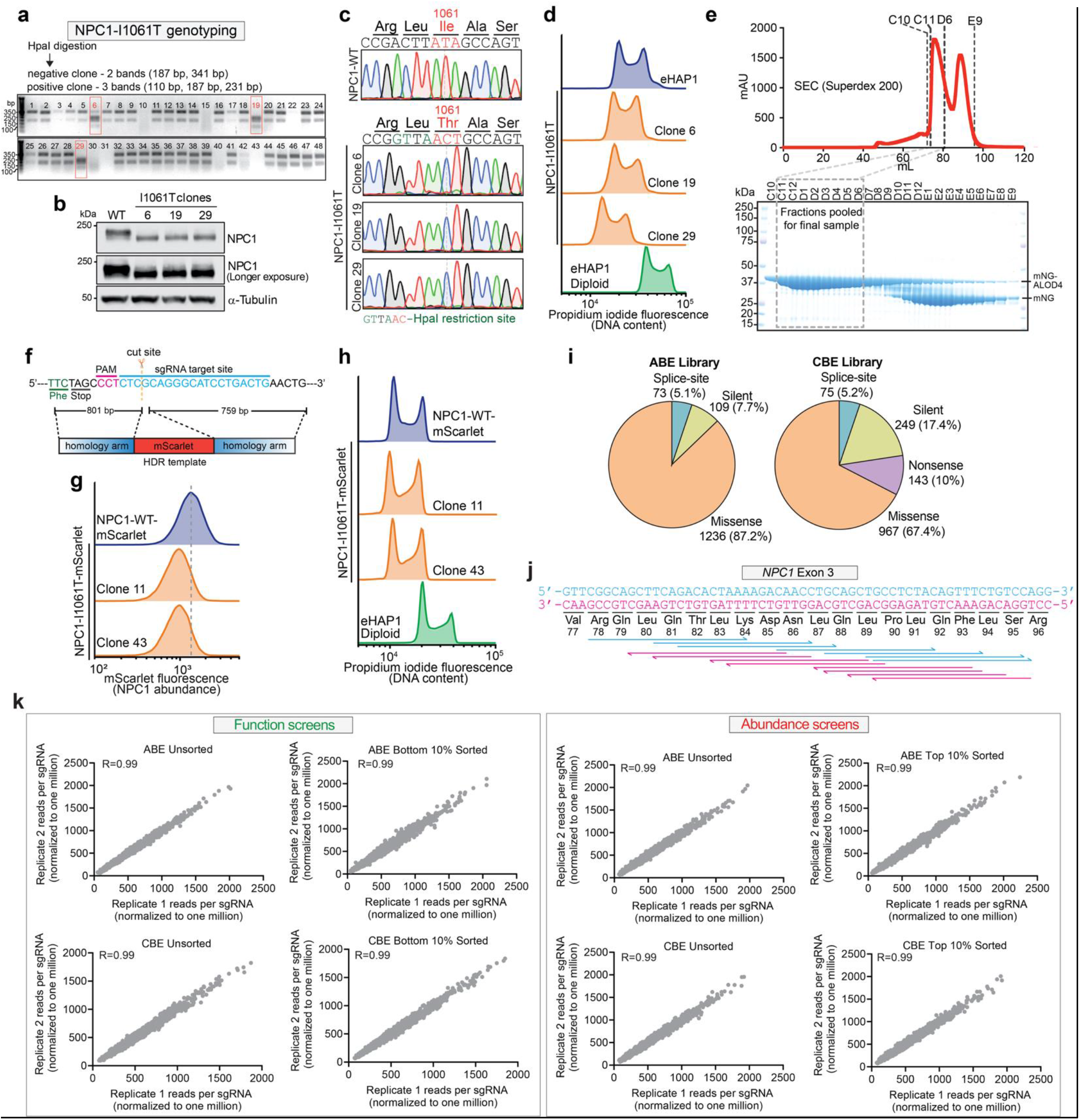
Haploid eHAP1 reporter cell lines and a tiled NPC1 base editing library for paired function and abundance screens. **a-d,** Homology-directed repair (HDR) was used to introduce the I1061T mutation and a linked HpaI restriction site at the *NPC1* locus in eHAP1 cells. Three clonal cell lines (#6, #19, #29) were chosen based on sensitivity to HpaI (**a**), immunoblotting to assess mutant protein abundance and SDS-PAGE mobility (**b**), and amplicon Sanger sequencing (**c**). Due to misfolding and ER-retention, NPC1-I1061T is known to be present at lower abundance and migrate faster on SDS-PAGE gels. Edited eHAP1 cell lines were confirmed to be haploid by ploidy analysis (**d**) prior to introduction of the base editing library. **e,** Size-exclusion chromatography profile of the mNeonGreen-ALOD4 (mNG-ALOD4) probe used to measure lysosomal cholesterol abundance by immunofluorescence and flow cytometry. The Coomassie-stained gel below shows the protein content of each fraction, with the mNG-ALOD4 band indicated. **f,** Schematic of the CRISPR/Cas9-assisted HDR strategy used to introduce a C-terminal mScarlet tag into the endogenous *NPC1* locus of eHAP1 cells. **g,** mScarlet fluorescence (a measure of NPC1 abundance) and **h,** ploidy analysis in two engineered eHAP1 NPC1-I1061T-mScarlet clonal cell lines. **i,** Predicted proportion of missense, silent, splice-site and nonsense mutations generated by the ABE and CBE libraries. Only CBE sgRNAs generate nonsense mutations. **j,** The tiling nature of the base editing library is illustrated using the *NPC1* Exon 3 region. Arrows below the sequence show the position of sgRNA sequences and are colored to match the targeted strand. Each codon in the exon is often targeted by multiple sgRNAs. **k,** Scatter plots showing the correlation of sgRNA read counts (per million total reads) between the two biological replicates for the unsorted populations and each of the four sorted populations using FACS gates described in **Fig.1d**.

**fig S2:**
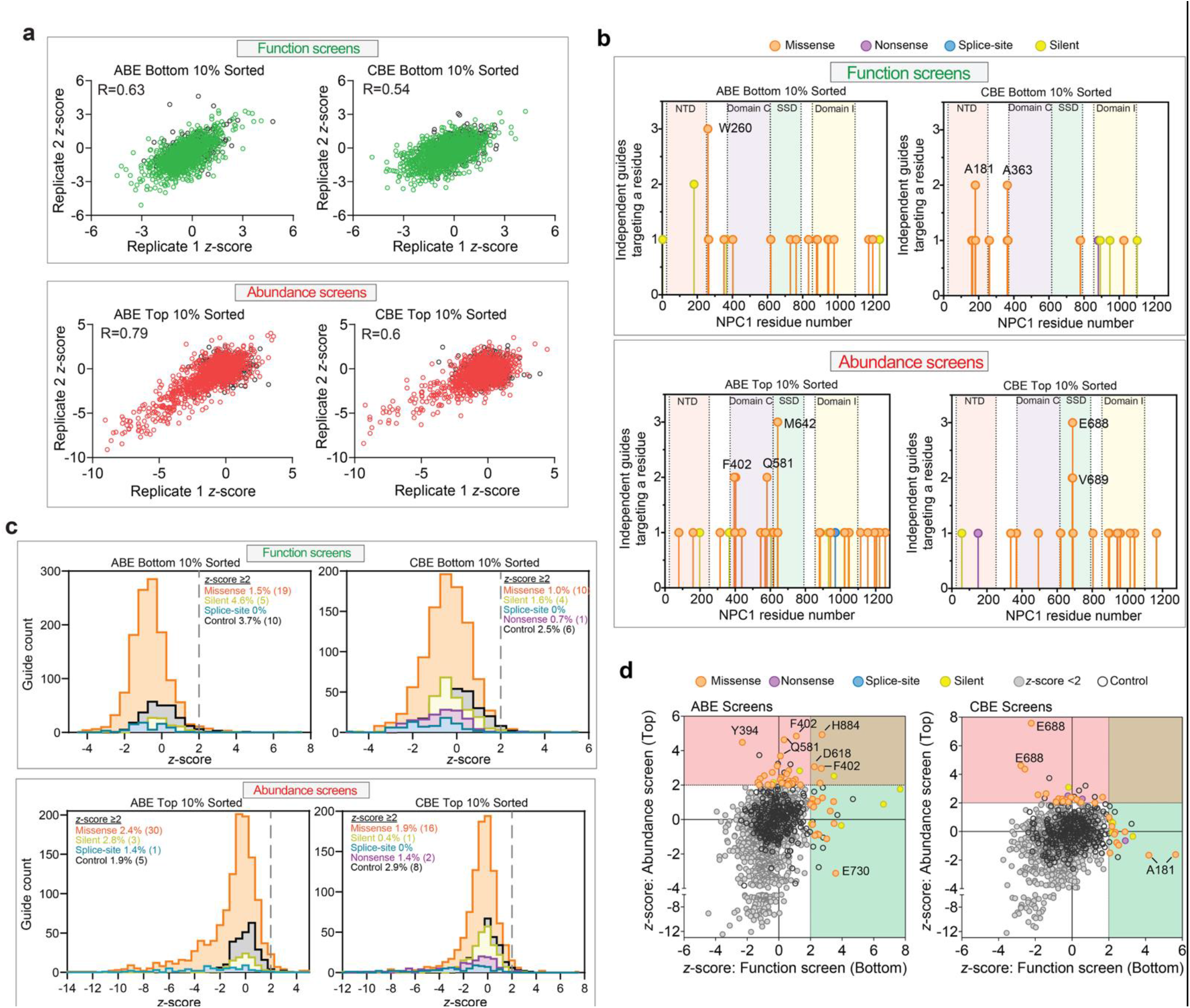
Reproducibility, sgRNA enrichment, and cross-modality analysis of the NPC1 base editing screens. **a**, Scatter plots showing the correlation of z-scores between the two biological replicates for each of the four screens diagrammed in **Fig.1d**. Non-targeting control sgRNAs are shown as black circles. **b,** Lollipop plots showing number of independent guides with z-score ≥2 targeting specific amino acid residues across the length of NPC1. Individual lollipops are colored according to the mutation type. When an sgRNA was predicted to generate both missense and nonsense mutations, only the nonsense mutation is plotted. **c,** Enrichment of NPC1 targeting and control sgRNAs in the sorted population with reduced lysosomal cholesterol (bottom 10% of mNG-ALOD4 fluorescence) or increased NPC1 abundance (top 10% of mScarlet fluorescence). The percentage of sgRNAs in each category with z-score ≥2 (dashed line) is indicated. **d,** Cross-correlation plot for all sgRNAs across the function and abundance screens. Each dot is a single sgRNA plotted based on its function z-score (Bottom 10% mNG-ALOD4 fluorescence, x-axis) and abundance z-score (Top 10% mScarlet fluorescence, y-axis). sgRNAs are colored according to predicted mutation type and non-targeting control guides are shown as black circles. The colored quadrants indicate sgRNAs enriched in function (green), abundance (red) or both (brown) screens.

**fig S3:**
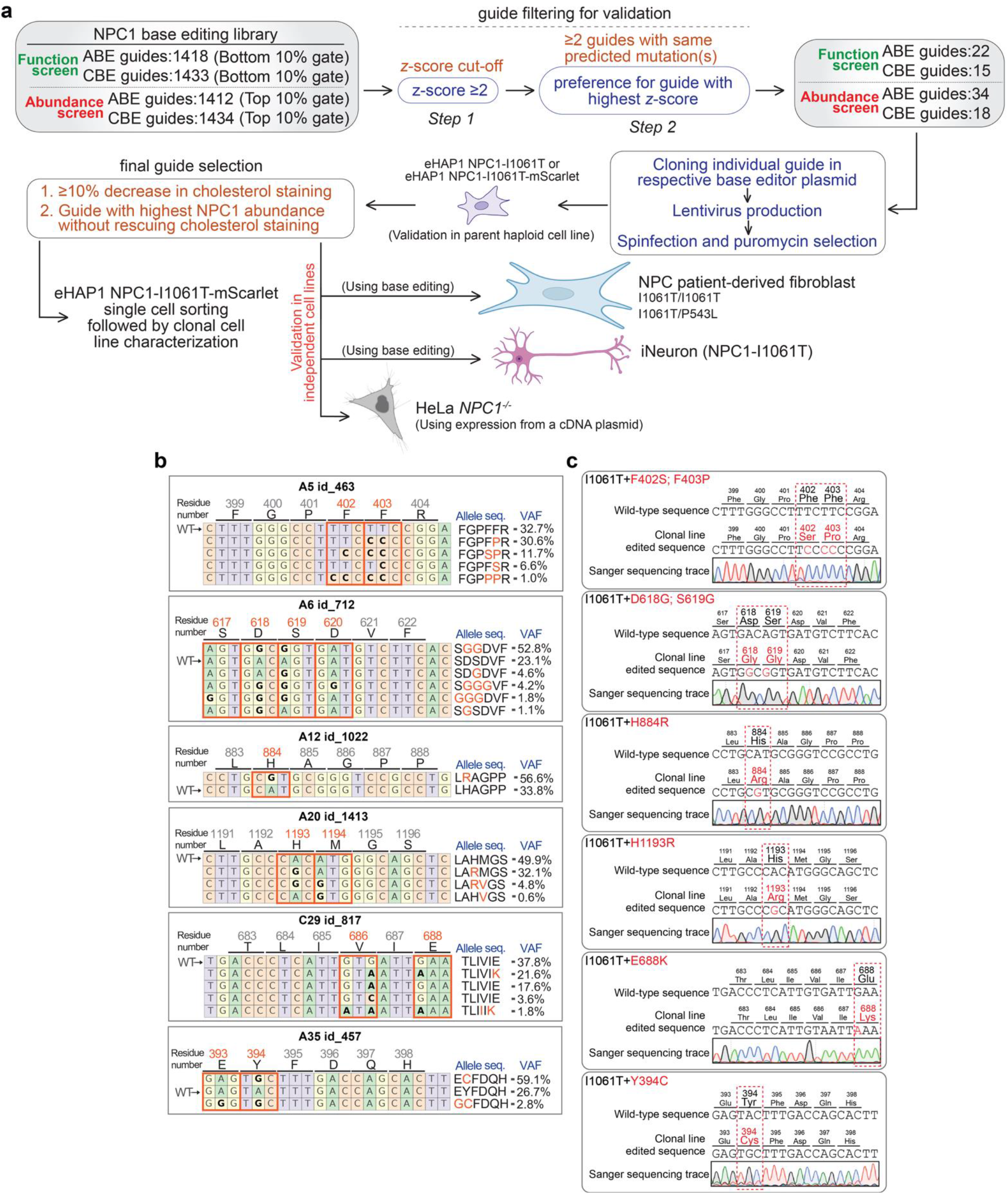
Validation pipeline and allele-level characterization of the six selected corrector sgRNAs. **a**, Outline of filtering and validation pipeline for sgRNAs targeting *NPC1* with z-scores ≥2. sgRNAs were individually introduced into eHAP1 NPC1-I1061T cells or eHAP1 NPC1-I1061T-mScarlet cells followed by assessment of lysosomal cholesterol clearance or NPC1 protein abundance. Selected sgRNAs were also validated in *NPC1^-/-^* HeLa cells, two different patient-derived fibroblast lines (NPC1*^I1061T/I1061T^* and NPC1*^I1061T/P543L^*) and iNeurons. **b**, Illumina sequencing (NGS) was used to measure variant allele frequency (VAF) in a pooled cell population after introducing 6 sgRNAs (A5, A6, A12, A20, C29, A35) (**Fig.2c**). Comparison of wild-type (black lettering) and edited (red lettering) sequence at each locus within the N3-N10 protospacer editing window shows the set of mutations produced by each sgRNA. **c,** Sanger sequencing chromatograms demonstrate the presence of second-site corrector mutations in NPC1-I1061T-mScarlet clonal cell lines used for mechanistic studies (**Fig.3**).

**fig S4:**
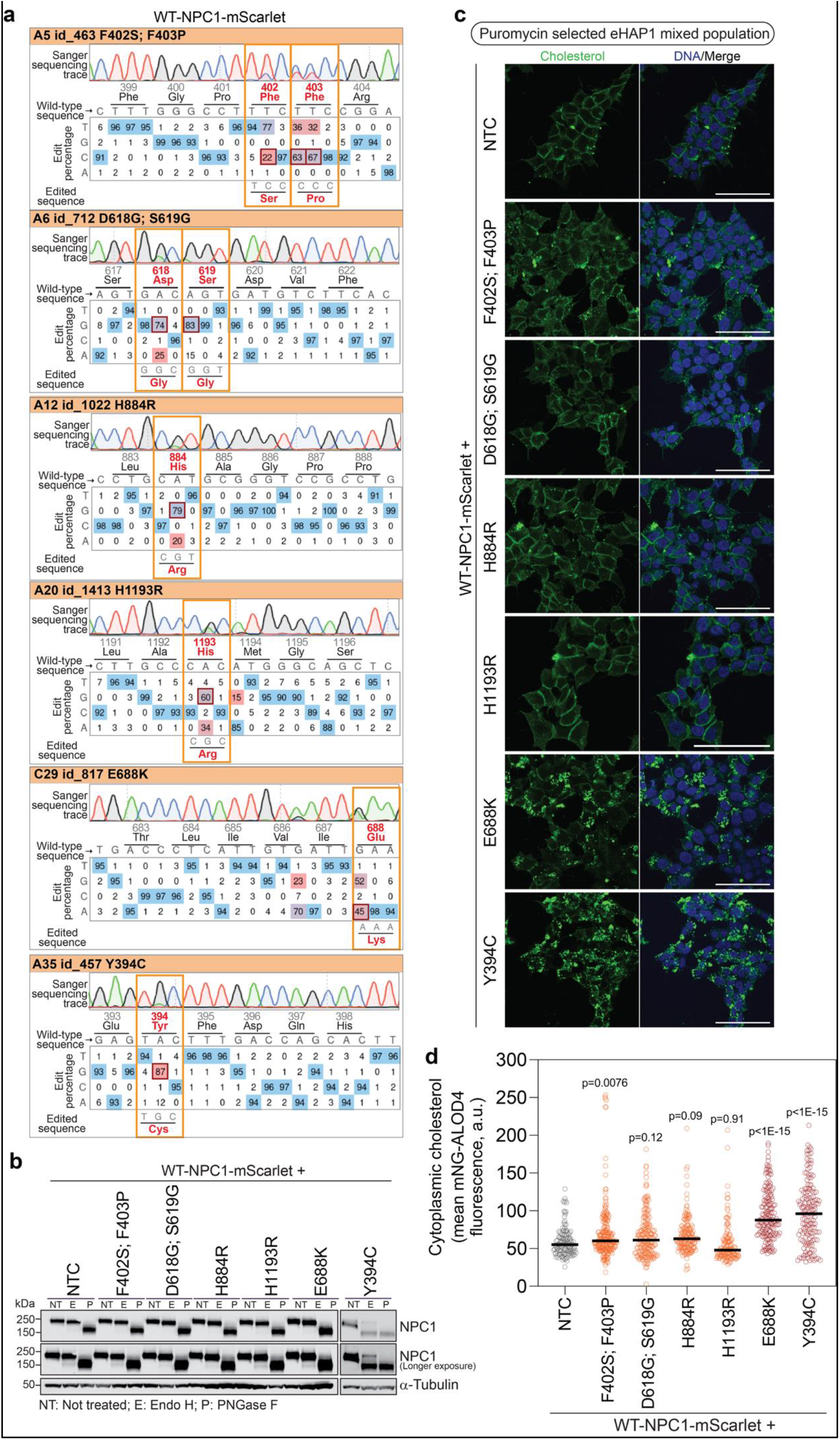
Effects of the six second-site mutations in wild-type NPC1 cells. **a**, Base editing outcomes in WT NPC1-mScarlet eHAP1 cells carrying the indicated sgRNAs (see **Fig.2c**). For each guide, Sanger sequencing traces are shown alongside wild-type and edited sequences, with edited codons (orange boxes) and edited nucleotides (red boxes) marked. Percentage of base editing was calculated using EditR (*51*). **b,** Glycosidase sensitivity assessed by NPC1 mobility on SDS–polyacrylamide gels (as in **Fig.3c**). Immunoblots showing fraction of NPC1 in ER (Endo H sensitive) or post-ER compartments (Endo H resistant). **c,** Lysosomal cholesterol was visualized in base-edited cells (shown in **a**) using mNG-ALOD4 staining. Nuclei are in blue. Scale bar: 50 µm. **d,** Quantification of cytoplasmic mNG-ALOD4 fluorescence from images in **c** was used to measure lysosomal cholesterol in each NPC1 variant (200-300 cells per variant, bar shows median, n=2). All experiments used pooled, not clonal, populations. Statistics: (**d**) one-way ANOVA with Dunnett’s post hoc test versus WT NPC1-mScarlet cells carrying a non-targeting control (NTC) sgRNA.

**fig S5:**
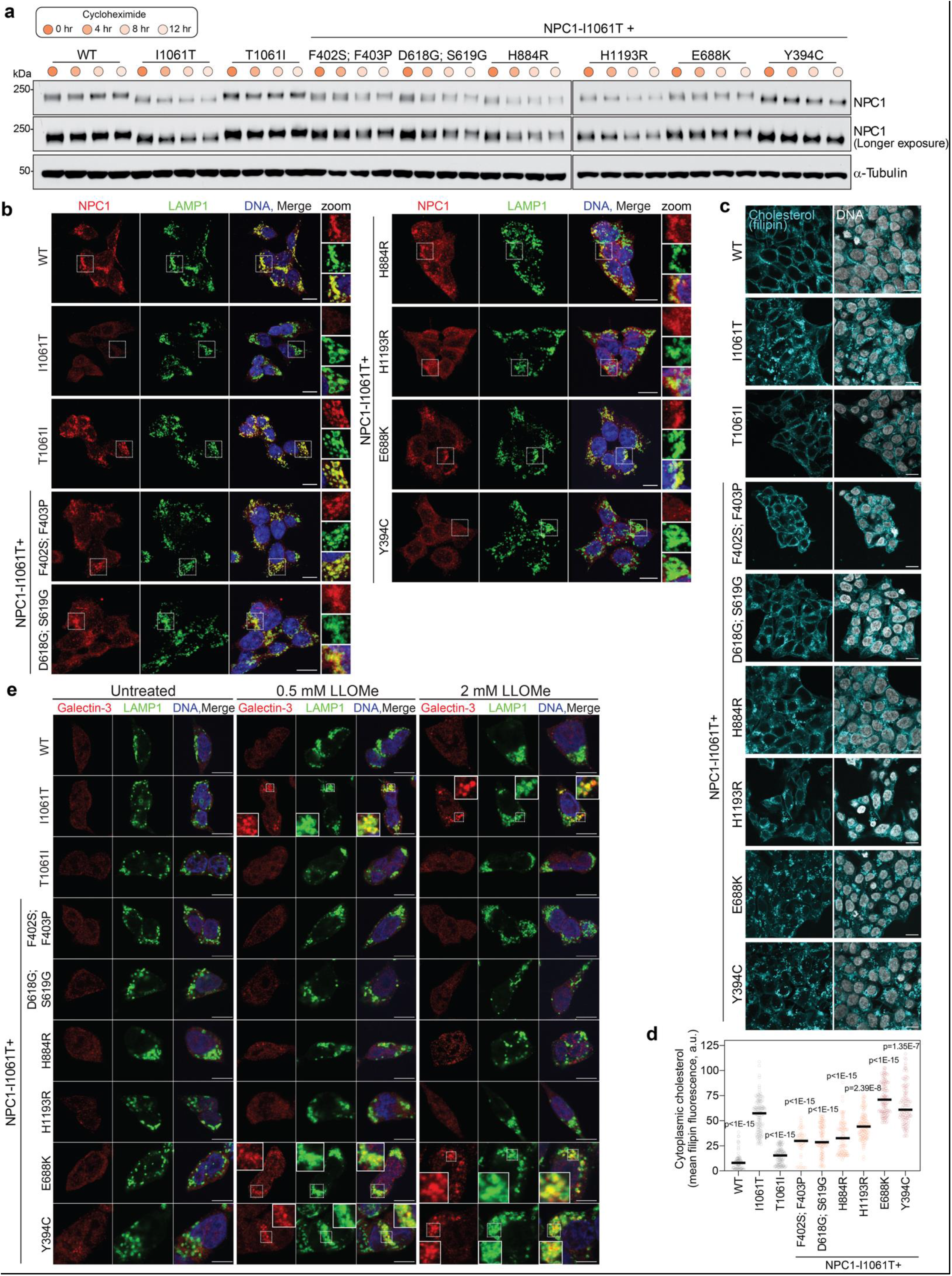
Cycloheximide chase blots, LAMP1 colocalization images, filipin staining, and LLOMe response images for the corrector variants from **Fig.3 a,** Representative blot showing the change in NPC1 abundance over time after addition of cycloheximide to block protein synthesis at time=0. α-Tubulin as loading control. Related to **Fig.3b**. **b,** Lysosomal localization of NPC1-I1061T variants (red) assessed based on their colocalization with the lysosome marker LAMP1 (green). Nuclei are in blue. Related to **Fig.3e**. **c,** Filipin staining was used to assess lysosomal cholesterol abundance in eHAP1 cell lines carrying the indicated variants of NPC1. Nuclei are in white. **d,** Quantification of cytoplasmic filipin fluorescence as shown in **c** (50-150 cells per variant, bar shows median). **e,** Assessment of lysosomal membrane integrity using Galectin-3 (red) recruitment following treatment with the lysosomotropic membrane-disrupting agent LLOMe. LAMP1 (green) was used to mark the lysosome. Nuclei are in blue. Related to **Fig.3h.** All scale bars in **b, c, e**: 10 µm. Statistics: (**d**) one-way ANOVA with Dunnett’s post hoc test versus I1061T.

**fig S6:**
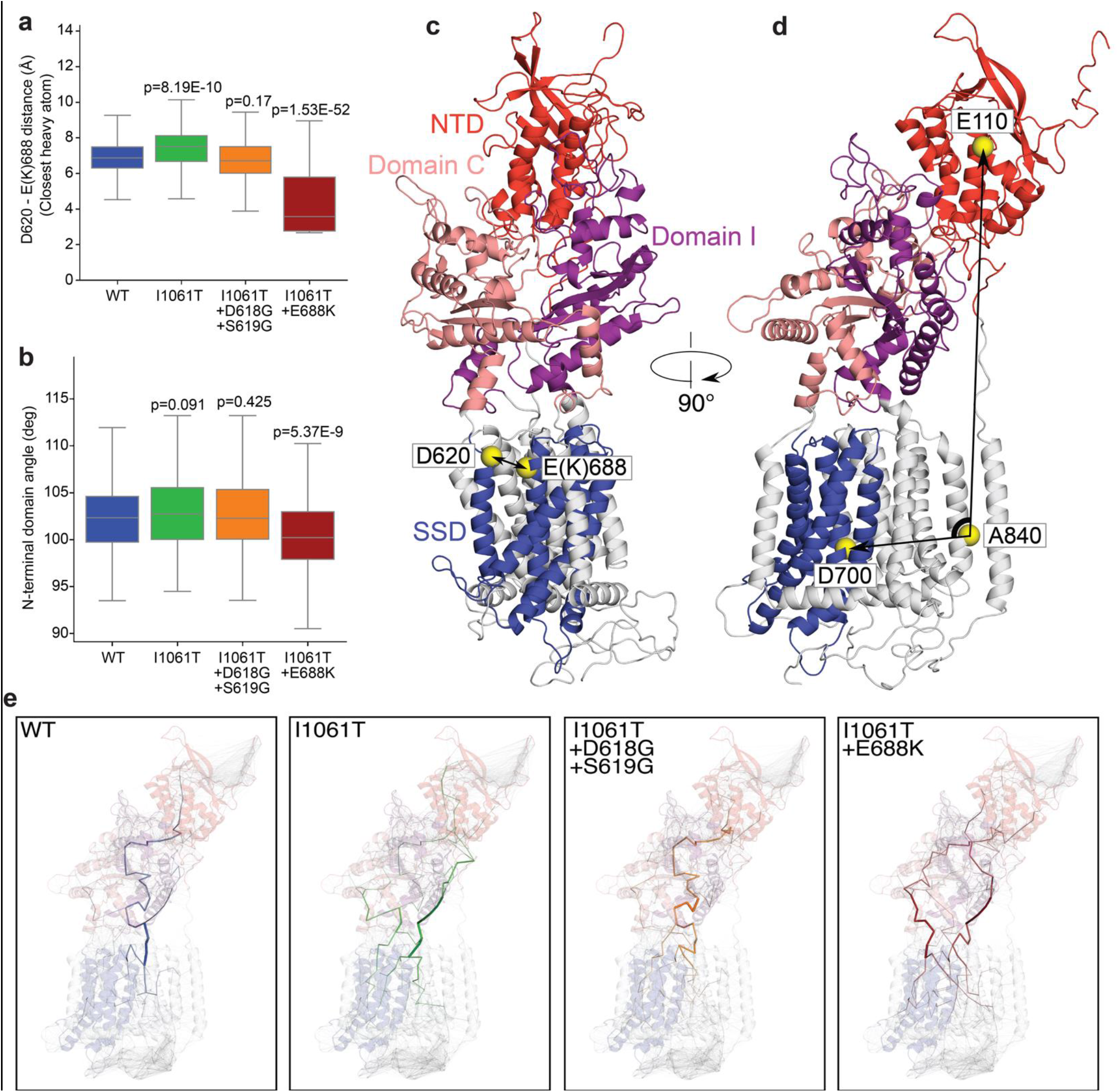
Molecular dynamics simulations reveal structural rescue mechanisms in NPC1 corrector variants. **a,c**, Distribution of pairwise closest heavy atom distances between D620 and residue E(K)688 (shown as gold spheres in **c**) in the indicated NPC1 variants. **b,d**, Distribution of the NTD–transmembrane angle (defined in **d**) for each NPC1 variant. **e,** Residue-residue interaction network analysis based on betweenness centrality of edges in contact networks derived from simulation trajectories. WT and D618G-S619G share similar network architecture with an intact information-transfer pathway between the NTD to the SSD, whereas the pathways are fragmented in NPC1-I1061T and NPC1-I1061T-E688K. Statistics: (**a,b**) Mann-Whitney U test was used to analyze probability distributions versus WT (300 trajectories per system).

**fig S7:**
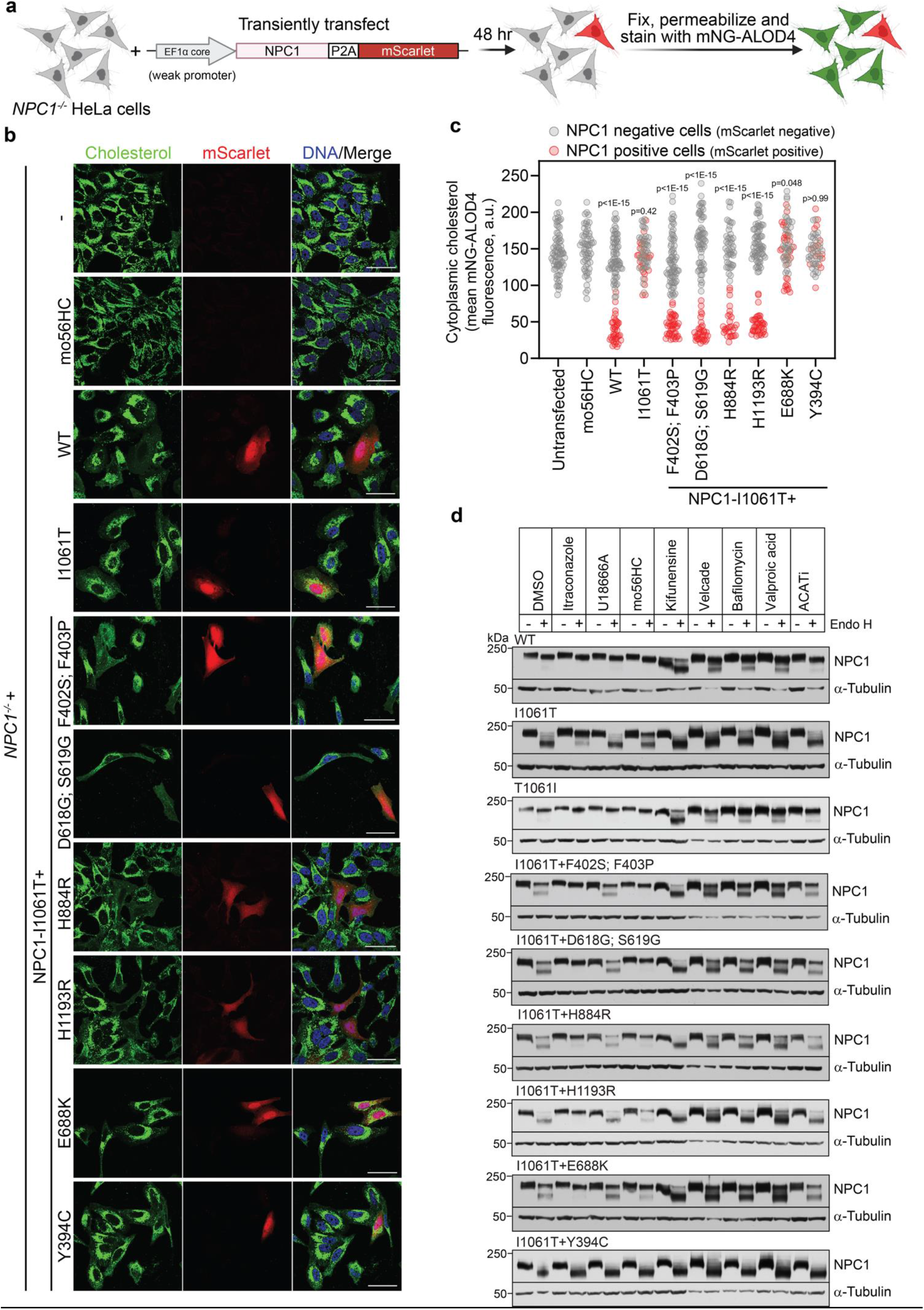
Orthogonal validation of corrector mutations by cDNA expression in *NPC1^-/-^* HeLa cells. **a**, NPC1 variants transiently expressed in *NPC1^-/-^* HeLa cells using core EF1α promoter to achieve low expression. A P2A-linked mScarlet reporter marked transfected cells. **b,** Correctors rescue lysosomal cholesterol accumulation in *NPC1^-/-^* HeLa cells, as evidenced by reduced mNG-ALOD4 fluorescence in mScarlet-positive transfected cells. Nuclei are in blue. Scale bar: 50 µm. **c,** Quantification of cytoplasmic mNG-ALOD4 fluorescence in mScarlet-positive (NPC1 expressing) and mScarlet-negative (NPC1 null) cells from the same population (50-150 cells per variant) indicates lysosomal cholesterol levels. **d,** Endo H resistance as a measure of post-ER pools of the indicated NPC1 variants treated with a panel of small molecules (raw data for heatmap shown in **Fig.5f**).

**fig S8:**
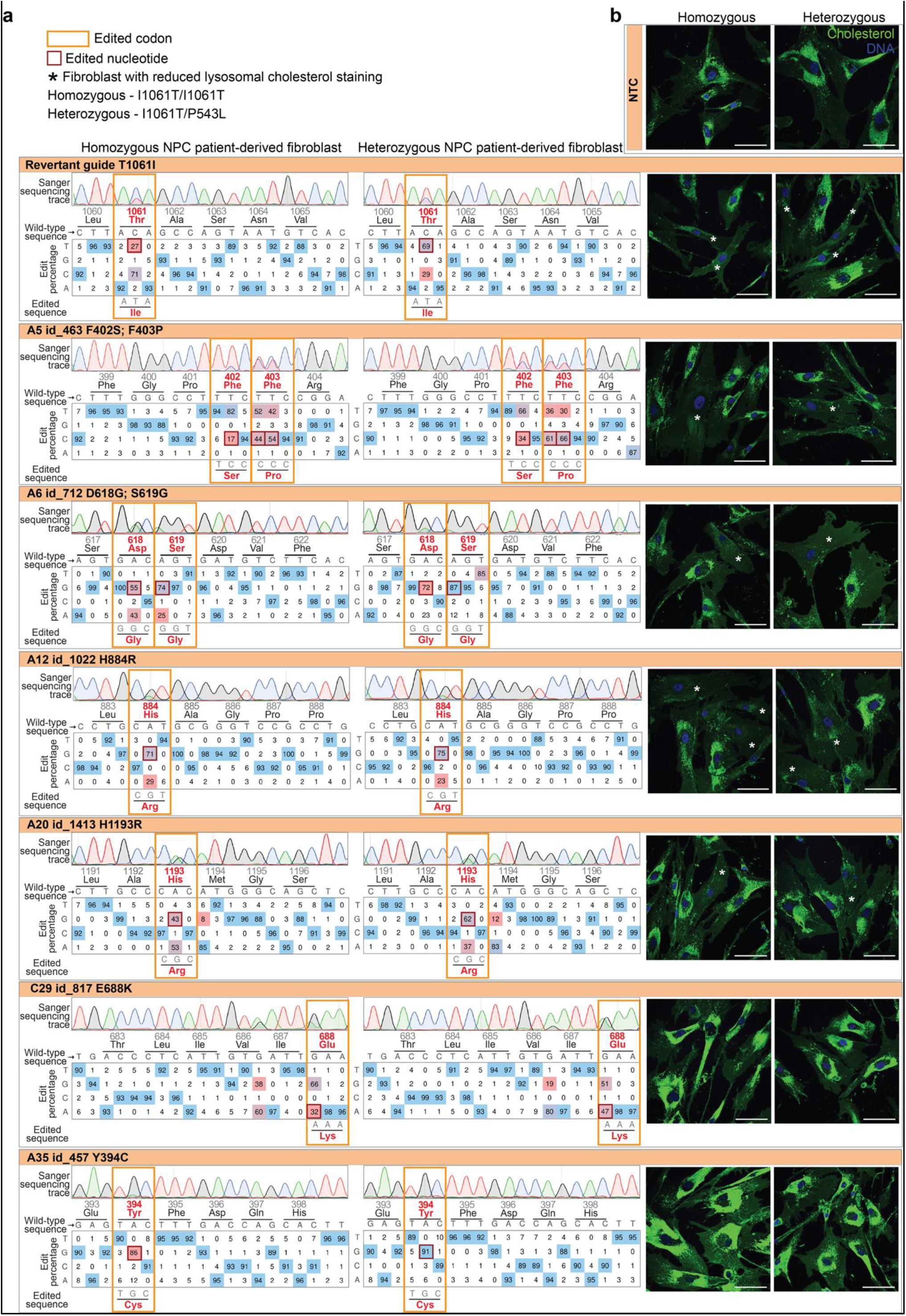
Base editing outcomes and cholesterol-staining images underlying the patient-derived fibroblast rescue data in Fig.6a. **a**, Base editing outcome analysis in homozygous (I1061T/I1061T) and compound heterozygous (I1061T/P543L) NPC patient-derived fibroblasts. For each guide, Sanger sequencing traces are shown alongside wild-type and edited sequences, with edited codons (orange boxes) and edited nucleotides (red boxes) marked. Percentage of base editing was calculated using EditR. **b,** Representative confocal images of NPC patient-derived fibroblasts stained with mNG-ALOD4 to detect lysosomal cholesterol accumulation after base editing using an NTC sgRNA (top) or NPC1 targeting sgRNAs that make the indicated second-site mutations. Asterisks (*) mark individual fibroblasts with reduced lysosomal cholesterol staining relative to surrounding unedited cells, indicating successful correction. Nuclei are in blue. Scale bar: 100 µm.

**fig S9:**
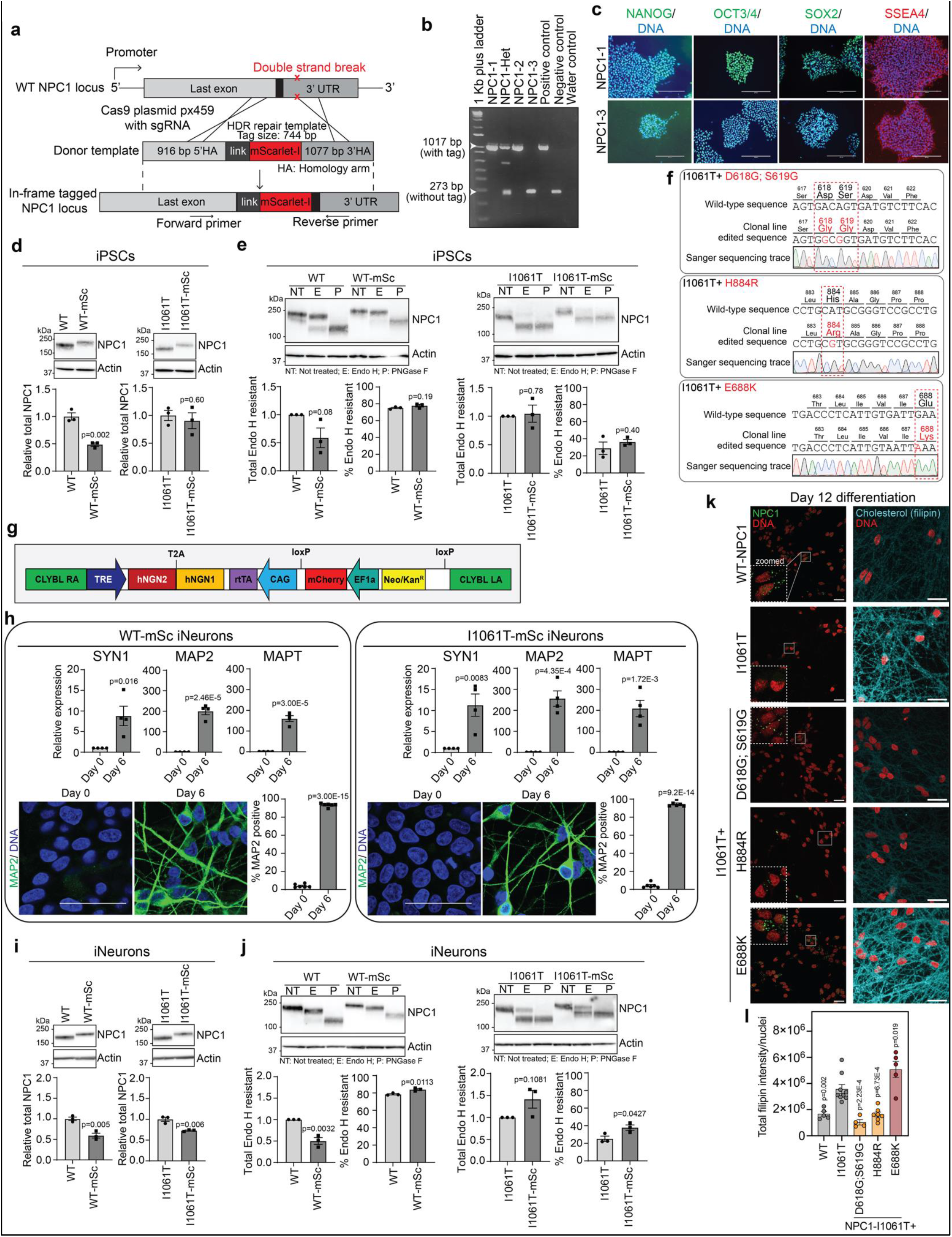
Generation and characterization of endogenously mScarlet-I–tagged NPC1 WT and I1061T iPSC and iNeuron lines. **a**, CRISPR/Cas9 knock-in of mScarlet-I at the endogenous *NPC1* C-terminus using a donor with 5′/3′ homology arms and sgRNAs targeting the terminal exon/3′ UTR. Genotyping primers outside the homology arms discriminate tagged from wild-type alleles. **b**, PCR genotyping of clonal iPSC lines: tagged allele = 1017 bp, untagged = 273 bp. Two homozygous clones (NPC1-1, NPC1-2), one heterozygous (NPC1-Het), unmodified NPC1-3, and controls shown. **c**, Selected clones retained pluripotency (NANOG, OCT3/4, SOX2, SSEA4 positive). Scale bar: 200 μm. NPC1-1 was chosen for downstream work based on low spontaneous differentiation and rapid proliferation. **d**, Immunoblot of total NPC1 in WT and NPC1-I1061T iPSC-derived cells ± mScarlet-I (mSc). Tagging significantly reduced NPC1 abundance in WT-mSc vs WT, but not in I1061T-mSc vs I1061T (mean ± SEM, n=3). **e**, Endo H sensitivity assay showed WT NPC1 was predominantly Endo H resistant (post-ER), whereas I1061T exhibited reduced resistance, consistent with impaired ER exit and lysosomal trafficking (mean ± SEM, n=3). **f,** Sanger sequencing chromatograms demonstrate the presence of second-site corrector mutations in I1061T-mSc clonal iPSC cell lines used for mechanistic studies. **g**, Doxycycline-inducible NGN2 cassette integrated into the CLYBL safe harbor locus via TALENs. **h**, Neuronal differentiation of WT-mSc and I1061T-mSc lines: qPCR showed significant induction of SYN1, MAP2, and MAPT after 6 days of dox *vs* undifferentiated. MAP2 immunofluorescence confirmed robust neurite outgrowth in both genotypes; MAP2⁺ quantification indicated efficient differentiation by day 6. Nuclei are in blue. Scale bar: 50 μm (n=3). **i**, Immunoblot of total NPC1 in WT and I1061T iNeurons ± mSc. Tagging significantly reduced NPC1 abundance in both backgrounds (mean ± SEM, n=3). **j**, Endo H sensitivity in iNeurons. In WT, mScarlet tagging reduced total Endo H resistant NPC1 but modestly increased %resistance, indicating preserved maturation of a smaller pool. In I1061T, tagging did not change total Endo H resistant NPC1 but significantly increased %resistance, suggesting partial rescue of mutant trafficking/stability post-ER during neuronal differentiation. WT remained predominantly mature; I1061T retained a large Endo H sensitive fraction (mean ± SEM, n=3). **k,** NPC1 abundance (left panels) and filipin staining (right panels) to assess cholesterol accumulation in a second set of iNeurons that were independently differentiated from iPSCs expressing the indicated variants of NPC1 with mScarlet tag. Nuclei are in red. Scale bar: 20 µm. **l,** Quantification of filipin fluorescence in iNeuron populations shown in **k** (50-100 cells per variant, mean ± SEM). Related to **Fig.6b-d.** Statistics: (**d,e,h,i,j**): unpaired *t*-test; (**l**) one-way ANOVA with Dunnett’s post hoc test versus I1061T.

**fig S10:**
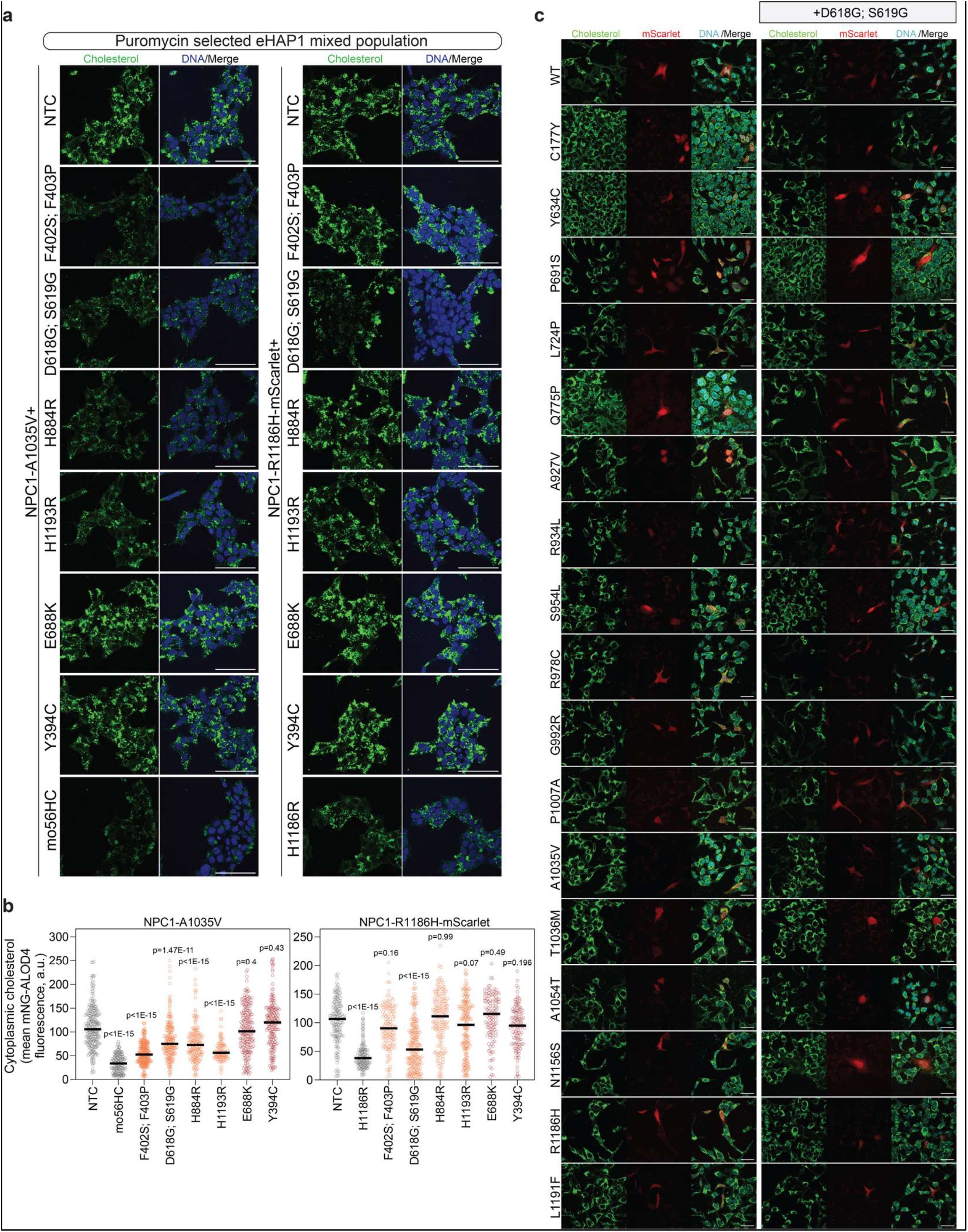
Comparison of corrector mutations on the NPC1-A1035V and NPC1-R1186H backgrounds and representative images for Fig.7b. **a,b**, Representative confocal images of eHAP1 NPC1-A1035V and NPC1-R1186H-mScarlet cells base edited with sgRNAs that introduce various corrector mutations. Lysosomal cholesterol was measured using cytoplasmic mNG-ALOD4 fluorescence and quantified in **b** (200-300 cells per variant, bar shows median). **c,** D618G-S619G corrector mutant rescues lysosomal cholesterol accumulation in *NPC1^-/-^* HeLa cells expressing diverse patient NPC1 variants, as evidenced by reduced mNG-ALOD4 fluorescence in mScarlet-positive transfected cells. Nuclei are in blue. All scale bars in **a,c**: 50 μm. Statistics: (**b**) one-way ANOVA with Dunnett’s post hoc test versus NTC.

## Notes

### Competing Interest Statement

The authors have declared no competing interest.

